# Visual physiology of the Layer 4 cortical circuit *in silico*

**DOI:** 10.1101/292839

**Authors:** Anton Arkhipov, Nathan W. Gouwens, Yazan N. Billeh, Sergey Gratiy, Ramakrishnan Iyer, Ziqiang Wei, Zihao Xu, Jim Berg, Michael Buice, Nicholas Cain, Nuno da Costa, Saskia de Vries, Daniel Denman, Severine Durand, David Feng, Tim Jarsky, Jerome Lecoq, Brian Lee, Lu Li, Stefan Mihalas, Gabriel K. Ocker, Shawn R. Olsen, R. Clay Reid, Gilberto Soler-Llavina, Staci A. Sorensen, Quanxin Wang, Jack Waters, Massimo Scanziani, Christof Koch

**Affiliations:** Allen Institute for Brain Science, Seattle, Washington 98109, USA; Janelia Research Campus, Howard Hughes Medical Institute, Ashburn, Virginia 20147, USA; University of California San Diego, La Jolla, CA 92093, USA; Novartis Institutes for BioMedical Research, Cambridge, MA 02139, USA; Howard Hughes Medical Institute and Department of Physiology, University of California San Francisco, San Francisco, California 94143, USA.

## Abstract

Despite advances in experimental techniques and accumulation of large datasets concerning the composition and properties of the cortex, quantitative modeling of cortical circuits under in-vivo-like conditions remains challenging. Here we report and publicly release a biophysically detailed circuit model of layer 4 in the mouse primary visual cortex, receiving thalamo-cortical visual inputs. The 45,000-neuron model was subjected to a battery of visual stimuli, and results were compared to published work and new in vivo experiments. Simulations reproduced a variety of observations, including effects of optogenetic perturbations. Critical to the agreement between responses in silico and in vivo were the rules of functional synaptic connectivity between neurons. Interestingly, after extreme simplification the model still performed satisfactorily on many measurements, although quantitative agreement with experiments suffered. These results emphasize the importance of functional rules of cortical wiring and enable a next generation of data-driven models of in vivo neural activity and computations.

**AUTHOR SUMMARY:** How can we capture the incredible complexity of brain circuits in quantitative models, and what can such models teach us about mechanisms underlying brain activity? To answer these questions, we set out to build extensive, bio-realistic models of brain circuitry employing systematic datasets on brain structure and function. Here we report the first modeling results of this project, focusing on the layer 4 of the primary visual cortex (V1) of the mouse. Our simulations reproduced a variety of experimental observations in a large battery of visual stimuli. The results elucidated circuit mechanisms determining patters of neuronal activity in layer 4 – in particular, the roles of feedforward thalamic inputs and specific patterns of intracortical connectivity in producing tuning of neuronal responses to the orientation of motion. Simplification of neuronal models led to specific deficiencies in reproducing experimental data, giving insights into how biological details contribute to various aspects of brain activity. To enable future development of more sophisticated models, we make the software code, the model, and simulation results publicly available.

## INTRODUCTION

Although our knowledge of the cortex has been improving dramatically thanks to the ongoing revolution in experimental neuroscience methods, the field is still far from an overall understanding of cortical circuits and their specific function. One essential component necessary to address this problem is the development of data-driven quantitative models that integrate experimental information and enable predictive simulations under a wide range of realistic *in-vivo-like* conditions – following the dictum attributed to Richard Feynman, “What I cannot create, I do not understand” [1]. Whereas detailed data-driven models of cortical tissue have been reported (e.g., [2]; see also [3]), modeling applications to an *in-vivo-like* regime have been limited (although see, e.g., [4] and references therein).

A typical systems neuroscience experiment involves a battery of different stimuli and, ideally, perturbations of the investigated circuit. Reproducing this in simulations of a data-constrained cortical model has proven challenging. To investigate the feasibility of *in-vivo-like* comprehensive simulations, and to build a platform for further studies, we set out to simulate a set of visual physiology experiments in the mouse primary visual cortex (area V1), with the emphasis on the first step in the cortical processing of visual information – namely, modeling the V1 input layer, the layer 4 (L4). We decided early on to replicate a small set of what we consider to be canonical physiological findings characterizing cells in L4 of V1. Given the thousands or more of published experiments carried out over the years in this region of cortex, our list is small, non-exclusive and may be considered idiosyncratic by some. However, we believe it is critical to start somewhere solid before generalizing indiscriminately. Our list of explored phenomena includes the approximately log-normal distributions of firing rates [5], orientation selectivity [6, 7, 8, 9], oscillatory population dynamics [10, 11, 12, 13, 14, 15], sparsity of responses to natural stimuli [16], amplification of thalamic inputs by recurrent connections [17, 18, 19, 20, 21, 22, 23], preferential connectivity among similarly tuned neurons [24, 25, 26, 27, 28, 29], and a number of others.

The model was constructed in a data-driven fashion from what is known about the L4 circuit organization. Although the model was biophysically detailed, we also used simplifications whenever possible, typically choosing computationally inexpensive approximations for biological mechanisms. A crucial component was a set of filters that represented visual information processing from image to the output of the lateral geniculate nucleus (LGN) of the thalamus, which projects to L4. This feature enabled one to use arbitrary movies as visual stimuli. We presented the same or similar sets of stimuli to the model and to mice in experiments and then compared the *in silico* and *in vivo* responses.

We asked three major questions: (1) How well does our model reproduce experimentally observed neural responses from the above list? (2) What are the major mechanisms that determine the neuronal activity patterns? And (3), how does the ability to reproduce experimental recordings depend on the level of granularity of the model?

To address (1), we assessed neuronal responses to artificial (e.g., drifting gratings) and naturalistic (e.g., movies) stimuli and selected a number of features of these responses that are generally considered important and interesting in the field. We then benchmarked the model performance on these features against the experimental data. Whereas typically models are developed to explain a specific phenomenon and may aim to reproduce 1-2 observed quantities, the key element in our study was to observe generalization over a wide variety of visual stimuli and response features. We found that our simulations reproduced many of experimental observations (with some exceptions) under a range of different stimuli.

For (2), we performed *in silico* experiments to investigate how individual neurons process inputs from different sources and how recurrent connections shape the network activity. This approach relied in part on simulated optogenetic experiments paralleling *in vivo* optogenetic studies. The most striking observation was that tuning properties of neurons were critically affected by the functional connectivity rules.

For question (3), we introduced two much simplified versions of our model, where biophysical neuron models were replaced by point-neuron models with either instantaneous or time-dependent course of synaptic action, and compared simulations of these simplified networks with the biophysical case. We found that, although the simplified network models qualitatively reproduced the trends observed in the biophysical simulations, the quantitative agreement with experiment suffered. The time-dependent synaptic kinetics in the simplified model allowed for better agreement with the biophysical model and experiment, such as, for example, in terms of producing oscillations in a gamma range.

No circuit in the brain exists in isolation, but including all the brain complexity in the model is impossible at present due to absence of data and inadequate computing resources. Therefore, we attempted to build a model of L4 of V1, which can be handled with available resources and for which a substantial amount of information could be found. The good performance of the model compared to the experiment may indicate a relative compartmentalization of L4 computation (see Discussion); this should not be expected everywhere in the cortex. Our L4 model provides a comprehensive characterization of activity and mechanisms in this cortical circuit and may serve as a stepping-stone for future, more sophisticated studies of all cortical layers. To enable this, we make the software code, the model, and simulation results publicly available (see SI).

## RESULTS

### Construction and optimization of the model

The network (**Fig. 1a, b**) consisted (see Methods) of models of individual neurons [30] from an early version of the Allen Cell Types Database [31], employing compartmental representation of somato-dendritic morphologies (∼100-200 compartments per cell) and 10 active conductances at the soma that enabled spiking and spike adaptation. Although recent additions to the Allen Cell Types Database include models of neurons with active conductances in the dendrites as well, those models are very computationally expensive, which was prohibitive for the breadth of our study (see below). In addition, in terms of somatic spike output, the current versions of such models do not exhibit much better performance than the models with active conductances restricted to the soma [31], and, thus, we used these latter, cheaper models. The cells were distributed uniformly in a cylinder ∼400 μm in radius, representing the central portion of V1, and 100 μm height with density of 200,000 mm^−3^ [32]. Five single neuron models represent five “types” of neurons – three major excitatory groups as determined by Cre-lines (Scnn1a, Rorb, Nr5a1, 85% of all cells) and two groups of parvalbumin-positive fast-spiking interneurons (15%, denoted as PV1 and PV2), which form the majority of interneurons in L4 [33]. All 10,000 biophysical cells were exact copies of these five models. The cell models correspond to regular-spiking excitatory cells and fast-spiking interneurons (PV+); whereas non-PV+ interneurons do exist in L4, they are a relative minority [33], and therefore were neglected for simplicity. Furthermore, 35,000 much simpler leaky-integrate-and-fire (LIF) neurons, with only two groups – excitatory and inhibitory – were placed around the biophysically detailed “core” to prevent boundary artefacts (see SI). The complete model accounted for over half of V1 L4 cells (45,000 out of ∼70,000). Below, we primarily focus on the properties of the biophysical core circuit.

**Figure 1.**
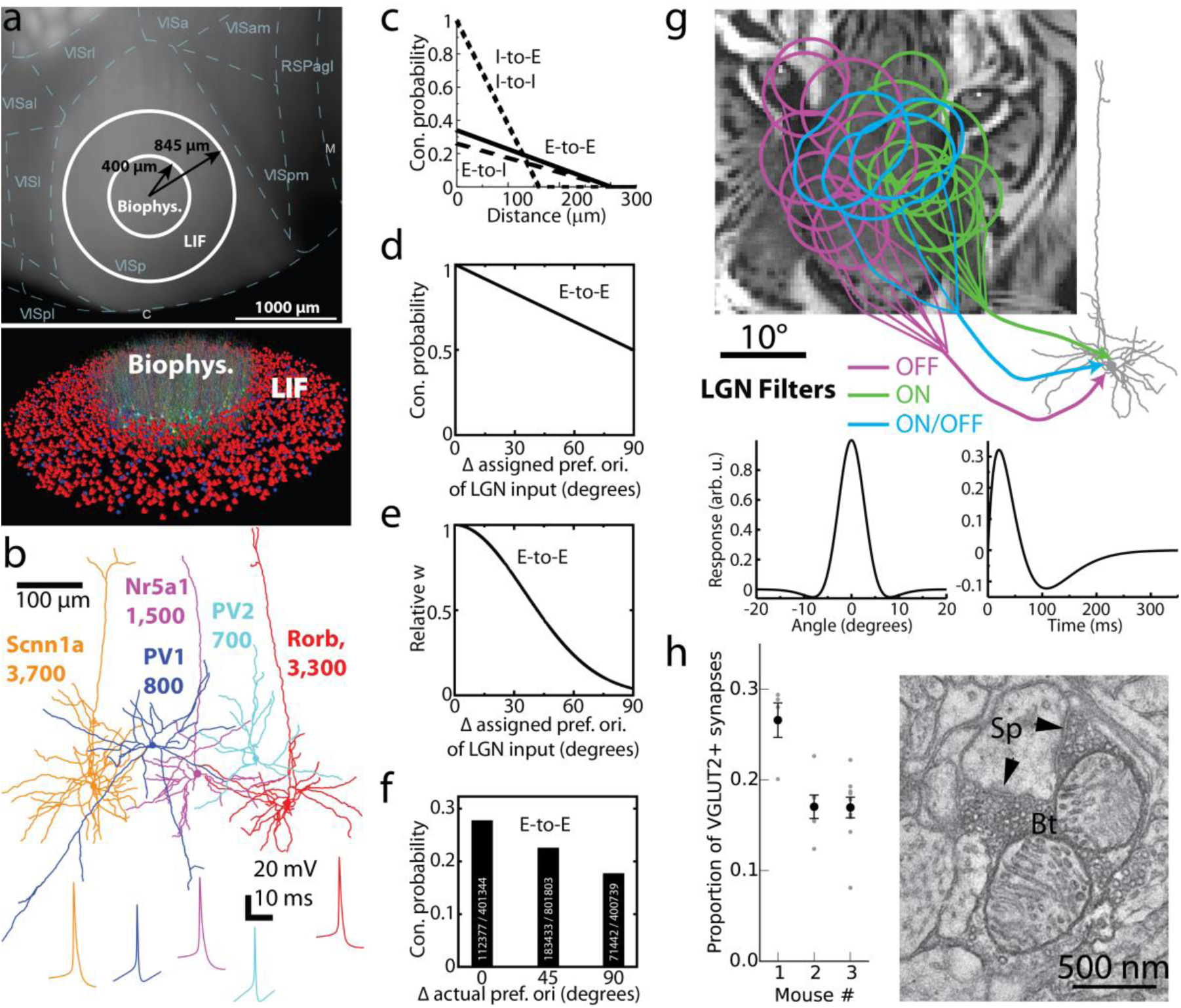
Construction of the model. (a) The biophysical and LIF portions of the model on the cortical surface with delineations of cortical areas (top; “VISp” is V1; the other labels starting with “VIS” correspond to higher visual areas) and with individual cells rendered (bottom; only a small subset of cells is shown for clarity). (b) Morphologies and action potential shapes of the five neuron models used to generate the L4 network; numbers of cells of each type are listed. (c-e) Connection probability (c,d) and synaptic weights (e) of excitatory (E) or inhibitory (I) cell targets and sources. The rules incorporate the dependence on the distance between somata (c) or difference of the assigned preferred orientations, *Δori* (d, e). Rules dependent on *Δori* were applied to E-to-E connections only, and synaptic weights for all connections were independent of distance. (f) Connection probability computed for cells within 50 μm using actual preferred orientation observed in simulations (cf. [24]). Numbers of connected and total pairs, used to obtain the probability, are shown inside the bars. (g) Three types of LGN filters (ON, OFF, and ON/OFF), superimposed onto an image, providing inputs to a L4 cell. Example filter’s spatial and temporal profiles are at the bottom. (h) The proportion of excitatory thalamocortical synapses (VGLUT2+) in the neuropil of V1 L4, as determined experimentally using EM. Proportions for individual samples of tissue are in gray; mean and s.e.m. for each mouse are in black. Right, an exemplar EM image of a putative VGLUT2+ synapse from the LGN onto an L4 neuron (Sp, spine; Bt, bouton; arrows: postsynaptic densities within the spine).

Three independent instantiations were generated using different random seeds. The connectivity and inputs into these three model instantiations were distinct, but all followed the rules described below. All simulations were performed using the python 2.7 code (an early version of the BioNet package [35]) employing NEURON 7.4 [34].

Recurrent connections (**Fig. 1c**) were established randomly according to probability decaying with intersomatic distance (e.g., [36]). Recent experiments [25, 26, 27, 29] showed that excitatory neurons in the mouse V1 L2/3 exhibit more likely and stronger connections if neurons prefer similar stimuli (“like-to-like connectivity”). We sought to investigate how such a rule affects neural activity. The rule (**Fig. 1d**) was applied to all excitatory cells by assigning a preferred orientation to each excitatory neuron (which was also used to select LGN inputs, see below) and using the difference between such orientations to compute a probability of connection. A similar like-to-like rule was applied to the amplitude of synaptic weights (**Fig. 1e**). The like-to-like rules were parameterized to correspond approximately to the observations for L2/3 [25] (**Fig. 1f**) and did not apply to pairs containing one or more inhibitory neurons. See Methods for further details.

We directly converted visual stimuli to spikes of LGN cells (which provide the input to L4) via linear space-time separable filters (**Fig. 1g**; [37, 38]) of ON, OFF, and ON/OFF types (see Methods), all of which produced relatively transient responses. Such a considerable simplification of the wide variety of response types that the LGN cells exhibit (see, e.g., [17]) was necessary to make the model tractable, especially because rules of connectivity from various functional LGN cell types and L4 in the mouse are largely unknown. Because the retinotopy of the complete model did not cover the entire field of view, we employed 9,000 LGN filters – approximately half of the mouse LGN. Based on the experimental data about receptive fields of L4 neurons [17, 32], we used retinotopy of L4 cells to “pool” inputs from LGN filters with similar retinotopy, and the assigned preferred orientation (see above) to establish the geometry of the ON and OFF subfields (**Fig. 1g**; see Methods, **Fig. 7**). This helped to establish orientation selectivity in the combined LGN inputs to individual L4 cells [32]. Interestingly, we found that in our model the L4 cells that were connected to each other had a higher chance to receive inputs from the same LGN cells, in comparison with unconnected L4 cells (see Methods, **Fig. 7d**). This is consistent with recent experimental observations for L4 and L2/3 cells [27].

The numbers of synapses for recurrent connections were chosen based on the literature (see Methods). Using electron microscopy (EM), we found (**Fig. 1h**) that LGN synapses constitute 15-30% of all synapses in the neuropil in L4 (see Methods for details), consistent with observations for mouse S1 and M1 [39]. This corresponds to over 1,000 synapses from the LGN per excitatory L4 neuron (given ∼8,000 synapses total per mouse V1 cell [32] on average, we assume relatively small L4 neurons to receive ∼6,000 synapses). Thus, the number of LGN-to-L4 synapses is an order of magnitude higher in the mouse than in the cat V1 (for which this number has been long established to be ∼100 [40]). These new data determined the numbers of LGN-to-L4 synapses in the model (see Methods for further details). For majority of connections, multiple synapses per connection were assigned, typically in the 3-7 range (see Methods). Synaptic dynamics was described by a sum of two decaying exponentials. We assumed no short-term plasticity as it is plausible that *in vivo* synapses operate at steady-state conditions [18]. Long-term plasticity was neglected because simulations covered several seconds at most.

To account for inputs from the rest of the brain and for different brain states, we introduced externally generated waves of background activity sweeping across the L4 model (**Fig. 2a**), inspired by reports of activity propagation over cortical surface *in vivo* [41]. Poisson spike generators were distributed within the cortical plane and were activated whenever the wave swept through their positions, sending spikes to the nearby L4 cells. This enabled the L4 model (**Fig. 2b**) to operate with an extremely simple analogue of distinct cortical states (see, e.g., [42, 43, 44]), which we call “Bkg. on” and “Bkg. off.” (see Methods). The random generation of the background waves, as well as random generation of background spike trains from the wave profiles and of LGN spike trains from deterministic filter outputs were the sole sources of indeterminacy, leading to differences between individual trials.

**Figure 2.**
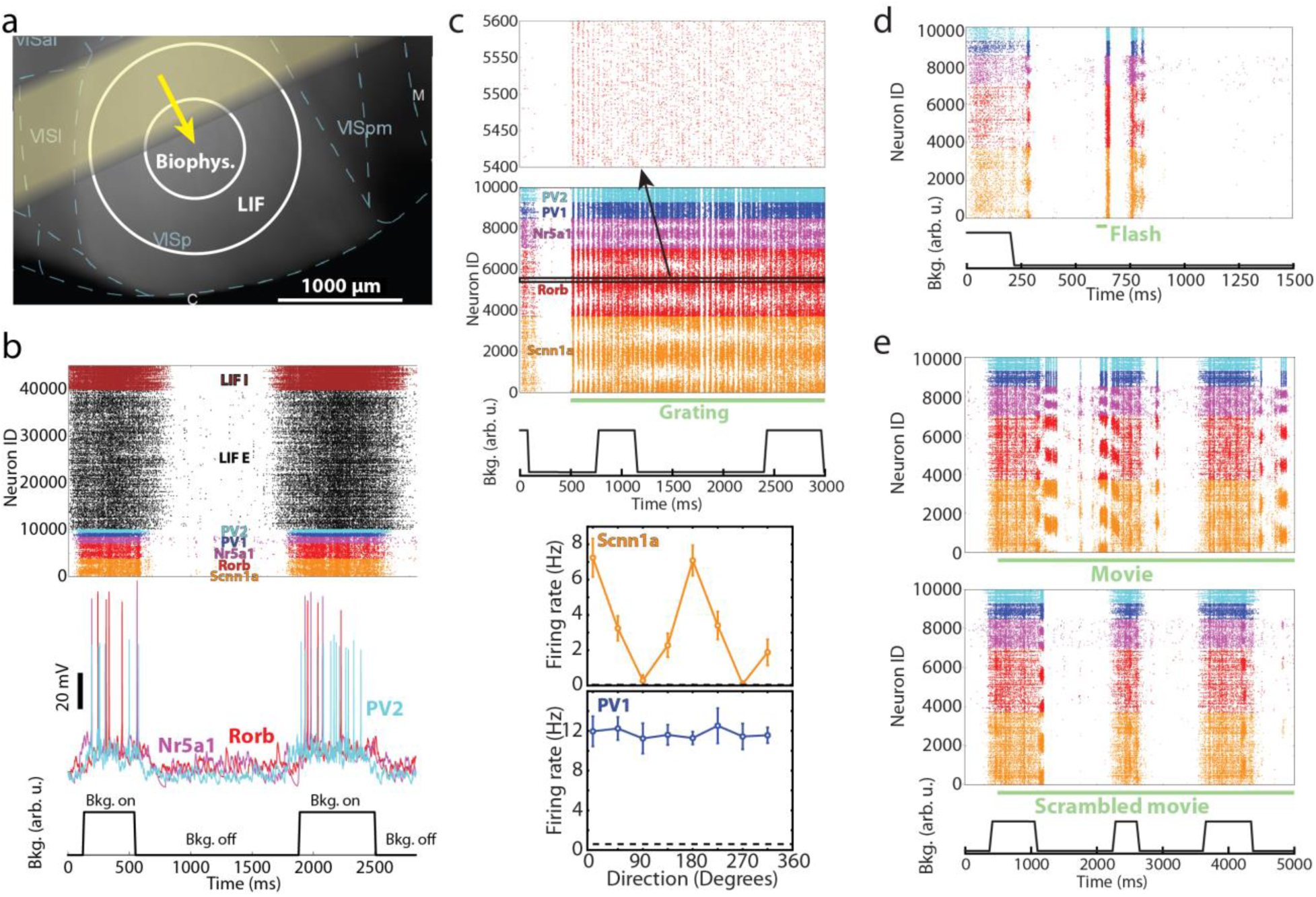
Examples of simulated responses to various visual stimuli. (a) Spontaneous activity in the model was generated by waves of background excitation (“Bkg. on”, yellow arrow denotes the direction of motion of the yellow bar-like region) alternated with intervals of no background excitation (“Bkg. off”). (b) Model activity in a gray screen trial. Examples of membrane potential traces from simulated biophysical cells are in the middle. (c) Spike raster in response to a drifting grating (TF = 4 Hz, at 0 degrees direction). Bottom, example orientation tuning curves for an excitatory and inhibitory cell from simulation. (d) Spikes in response to a 50 ms full-field flash. (e) Spike raster for a single trial of a natural movie (top) and for a temporally scrambled version of the same movie (bottom). The raster in (b) shows all neurons and those in (c-e) for clarity show the 10,000 cells in the biophysical core of the model (inset on top of (c) zooms in on 200 cells). All rasters are examples from one trial; all trials used unique combinations of “Bkg. on” and “Bkg. off” states (shown at the bottom of plots), which overlapped (or not) in different ways with the visual stimuli.

After the steps above, all parameters were fixed except the synaptic weights (conductance amplitudes). The LGN-to-L4 weights were selected to produce experimentally observed excitatory currents from LGN [17]. The weights for recurrent connections were then manually optimized, while constraining post-synaptic potentials and currents (PSPs and PSCs) to the experimentally reported range for mouse cortex [45] (see Methods). Optimization was limited to two training stimuli – a single trial of a drifting grating and one of a gray screen, 500 ms each. The target was to reproduce the mean spontaneous firing rate and the rate in response to the preferred grating (*R*_*max*_), while avoiding synchronous epileptic-like activity. Published data from [N6] were used for this optimization, before our own experiments were finalized. After optimizing the synaptic weights using this limited training set, we applied the model without any further modification to all the other stimuli, which served as a test set. The exact synaptic weights obtained during the optimization stage may not represent a unique solution, due to degeneracy [46]. Nevertheless, it is reassuring that synaptic properties from our optimized models were consistent with published experimental reports (see **Fig. S2**), exhibiting, e.g., current amplitudes of ∼10-30 pA for excitatory and ∼40-60 pA for inhibitory synapses [45] or voltage amplitudes of ∼1 mV [47] for L4 excitatory-to-excitatory connections (as measured at the soma).

### Benchmarking simulated neuronal activity to the experimental data

How well does the model reproduce experimentally observed neural responses *in vivo?* The answer will depend on the types of stimuli and features of responses to those stimuli that one chooses for comparison. We reasoned that a successful model should be versatile; in other words, it is not sufficient to be able to model responses to one type of stimuli well, and instead one should strive to reproduce features across many types of stimuli. Therefore, our battery of visual stimuli included gray screen, a variety of drifting gratings, 10 natural images, 3 natural movie clips, 2 types of full-field flashes, and 4 moving-bar stimuli (see example responses in **Fig. 2c-e**; see also Methods, **Fig. S3**, and **Table S2**). Altogether, ∼3,600 simulations were carried out. Our own experiments used for benchmarking consisted of two series of extracellular electrophysiological recordings in mouse V1 employing multi-electrode silicon probes. In the first [7], responses to drifting gratings with a variety of spatial and temporal frequencies were measured, in both awake and anesthetized mice. In the second (previously unpublished), spontaneous activity (responses during gray screen presentation) as well as responses to natural movies, natural images, full-filed flashes, and other stimuli were recorded in awake mice. We chose several features of visual responses that are generally considered important and are often reported in the field of cortical visual physiology and compared their values between experiment and simulations (**Fig. 3**). It should be noted that, although the measured metrics differed slightly among the three excitatory cell types (Scnn1a, Rorb, and Nr5a1), as well as among the two inhibitory types (PV1 and PV2), we do not ascribe significance to these differences. This is because during model construction and optimization no experimental data was available on distinctions in response features or connectivity between these L4 cell populations, and, in absence of either, we assumed the same optimization targets for all excitatory or inhibitory cells (which, however, allowed for ∼1 Hz rate deviation from the target, resulting in different types settling at slightly different activity levels).

**Figure 3.**
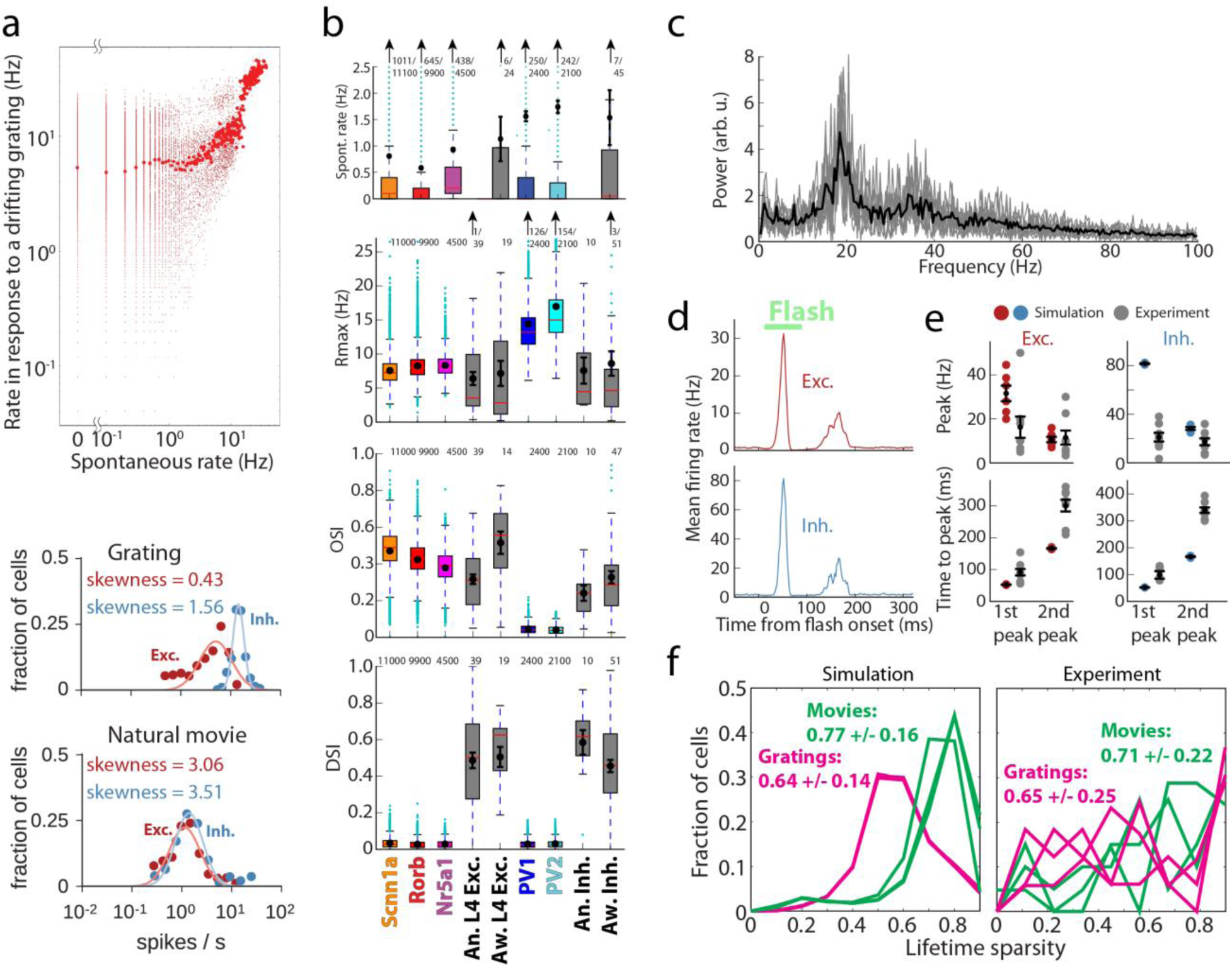
Benchmarking the simulation results. (a) Log-normal like distribution of firing rates. Top, firing rates of all biophysical cells from three models during spontaneous activity and in response to a drifting grating, averaged over 10 trials. Large red dots are firing rates averaged over all cells in 0.1 Hz bins over the spontaneous activity axis. Bottom, examples of firing rate distributions on a log scale for a single trial. Solid lines indicate log-normal fits of the data. (b) Comparison of the spontaneous rates, maximal rates in response to gratings (*R*_*max*_), orientation selectivity index (OSI), and direction selectivity index (DSI) between the simulation (by cell type, color) and experimental measurements using extracellular electrophysiology (gray). See Methods for details on the elements of the box plots shown here and in other figures. “An.” – experiments in anesthetized mice, “Aw.” – in awake mice. (c) The local field potential (LFP; see Methods) measured at the center of the L4 model, for a drifting grating. The spectra from 10 trials are shown in gray, and the averaged spectrum is in black. (d) The model PSTH in response to a 50 ms flash (average over all biophysical excitatory cells, all models, and all trials, in 2 ms bins). (e) The magnitude and time-to-peak (from flash onset) for the first and second peaks of the response to the 50 ms flash, for both simulation and electrophysiological data. (f) Distributions of lifetime sparsity of simulated and experimental responses of excitatory neurons, computed for three directions of a grating (0, 45, and 90 degrees) and for three movies. See Methods for details on computing all values presented.

The first characteristic we checked is whether neuron firing rates follow a positively-skewed, log-normal-like distribution, which is a ubiquitous hallmark feature of brain activity *in vivo* [5]. Such distributions were indeed observed in our simulations for spontaneous activity and all types of visually-driven responses (**Fig. 3a, S5a**). Consistent with published reports, our simulated rate distributions spanned 2−3 decades and may widen further in longer simulations (for instance, spontaneous rates were computed from 20 trials of 500 ms each, resulting in the lowest possible rate of 0.1 Hz). Also in agreement with literature [5], individual neurons tended to keep their low or high firing rates across different stimulus conditions (**Fig. 3a**, top). The positive skewness of firing rate distribution (i.e., the third standardized moment, 〈((*f* − 〈*f*〉)/*σ*_*f*_)^3^〉, where *f* is the firing rate and *σ*_*f*_ is its standard deviation, both computed in the linear (i.e., not log) firing-rate space) was typically between 0 and 4 (**Fig. S5a**). While it is hard to expect an exact match of the experimental skewness distributions from our relatively rough model, it is reassuring that the model reproduced the overall experimental trends of firing rate distributions with positive skewness in the 0-4 range. Note that a normal distribution has zero skewness, whereas log-normal distribution has positive skewness.

The mechanisms underlying log-normal like distributions of activity of individual neurons *in vivo* are not well understood [5], and may involve both cell-intrinsic and network phenomena. Our model does not clarify the potential contribution of the former, since all cell types were tuned to the same target rates (apart from the excitatory/inhibitory distinction), but sheds light on the latter. We found that, when similar levels of background excitation were applied to all cells, spontaneous rates were either zero or uniformly close to the mean, whereas log-normally distributed strength of background inputs (see **Fig. S1c**) resulted in spontaneous firing rates distributed over 2 or more decades. In a highly simplified approximation, one may consider that under visual stimulation cells preferring a given stimulus may increase their firing by about an order of magnitude, whereas activity of all other cells remains more or less the same as under no stimulation. Thus, given log-normal distribution of spontaneous rates, one expects that the distribution under visual stimulation remains largely the same, perhaps slightly wider, which is indeed what we observe (**Fig. 3a**). This emphasizes the importance of enabling log-normal like distribution of spontaneous rates, and our work suggests that log-normally distributed strength of inputs that cells receive (potentially, from non-local sources) may play a big role in this, although other mechanisms cannot be excluded.

Another aspect of activity that we checked was whether the magnitude of responses was consistent with the experiment; presumably, it is important for network computations that cells of certain types fire at certain rates in response to particular stimuli. For this purpose, we considered the rates of spontaneous activity and maximal rates (*R*_*max*_) in response to drifting gratings (**Fig. 3b**; see Methods for definitions). The spontaneous rates were 0.5-1.0 Hz for excitatory and 1.5-2.0 Hz for inhibitory cells, broadly consistent with our experimental measurements and published data [N6], and *R*_*max*_ levels were also similar to experimental ones.

Furthermore, the L4 responses to gratings are known to exhibit orientation and direction selectivity (e.g., [6, 7]). We found that the orientation selectivity index (OSI; see Methods) for excitatory cells was ∼0.4-0.5 on average, in excellent agreement with the experiments (**Fig. 3b**). For inhibitory cells, OSI was ∼0.05 on average, whereas the experimental value was ∼0.3. The relatively low OSI values were due to non-selective thalamic inputs and recurrent connectivity for inhibitory cells (see Methods), which were chosen that way at model construction to conform to an often-expressed view that excitatory cells are tuned and inhibitory fast-spiking cells are not. However, recent experiments [33], as well as our own data (**Fig. 3b**), suggest moderate levels of tuning for these neurons, and therefore future models will need to reflect these observations better. Nevertheless, qualitatively the model already captures the trend of better tuning of excitatory cells compared to inhibitory fast-spiking cells. Another much studied aspect of orientation tuning in the cortex is its contrast invariance in excitatory neurons (see, e.g., [8, 9]). Excitatory neurons in our simulations were strongly orientation-tuned for contrast as low as 10% (**Fig. S5c**). We computed the differences of the half-width at half-height of the tuning curves at high and low contrasts for each cell (ΔHWHH) and found (**Fig. S5d**) that the distribution of ΔHWHH sharply peaked near zero. On average, the difference was 4 +/−5 degrees for 80% vs. 10% and −2 +/−4 degrees for 80% vs. 30% contrasts (i.e., the average was within the standard deviation in both cases). Thus, our model exhibited substantial robustness of tuning curves to contrast (although perhaps not to the full extent of the real mouse V1 [8]).

By contrast to OSI, the model performed poorly on the direction selectivity index (DSI; **Fig. 3b**), which was close to zero in simulations, whereas experimentally it was ∼0.4-0.5. This was due to the extreme simplicity of the LGN inputs, which were constructed based on the experimental data, but at this point did not include all types of LGN activity observed experimentally [7, 17, 21]. In particular, for simplicity, we did not include the sustained LGN responses (i.e., all LGN filters produced transient responses), whereas the most recent experimental data suggest a critical role for the interplay between sustained and transient LGN inputs in generating direction selectivity in V1 [23]. Proper incorporation of direction selectivity is the subject of ongoing work on the next generation of the model.

Another important aspect of neuronal activity *in vivo* is the population-level oscillatory rhythms, which are observed in a variety of frequency bands. Whereas oscillations at many of such frequencies are likely caused by non-local interactions in the brain, and therefore cannot be expected to arise in an isolated L4 model, some of them may be generated locally. Indeed, our simulations exhibited oscillations in the 15-50 Hz range (sometimes referred to as“mouse gamma”), with a peak at ∼20 Hz (**Fig. 3c**). This is consistent with extracellular electrophysiology data [10], which exhibits a particularly strong peak at similar frequencies in L4.

Global luminance changes present another challenge to models, due to simultaneous engagement of all cells by these stimuli. We studied responses to 50 ms full-field flashes and observed, in both experiment and simulations, that a sharp and fast peak in activity was evoked by the first white-to-gray transition, followed by a second smaller and wider peak (**Figs. 2d, 3d**). The magnitude of the first peak was about 2-4 times higher in simulation than in the experiment, whereas that of the second peak was approximately the same, and the time course of the response was uniformly ∼2-fold faster in simulations (**Fig. 3d, e** and **Fig. S5b**). These differences were most likely again caused by the absence of sustained LGN responses in the model. Importantly, however, in both simulations and experiments, the second peak was delayed – instead of 50 ms (the flash duration), it appeared 100 ms after the first (200 ms in the experiment). This reflects a known phenomenon of suppression following luminance change, where the second peak corresponds to release from inhibition (e.g., [48, 49]). Thus, qualitatively the model reproduces well the critical features of cortical responses to global luminance change, which likely affect perception of visual stimuli (such as in, e.g., luminance-induced visual masking).

Finally, an essential observation is that natural scenes evoke responses in V1 that are substantially different from those evoked by artificial stimuli such as gratings. This was the case in simulations – for example, responses to a natural movie in **Fig. 2e** were very distinct from those to a horizontally drifting grating in **Fig. 2c** (both use the same ID labels). For a grating, cells with certain IDs respond strongly throughout the simulation (as they prefer the grating’s orientation). By contrast, natural movies evoked episodes of concerted responses in distinct populations of cells (which share similar orientation preference). Notably, such concerted responses were mostly absent when movie frames were shuffled in time (**Fig. 2e**, bottom). To quantify the differences, we computed lifetime sparsity for each cell following Vinje and Gallant [50] (see Methods). Sparsity was higher for movies than for gratings (**Fig. 3f**) in both simulations and experiments, consistent with previous observations (including Ca^2+^ imaging [16]): 0.77+/−0.16 in simulation vs. 0.71+/−0.22 in experiments for movies and 0.64+/−0.14 vs. 0.65+/−0.25 for gratings.

### Mechanisms underlying neural activity and computation in the L4 circuit

We further used the model to shed light on the mechanisms that determine patterns of neural activity and computations performed by the L4 circuit. One important question is to what extent the L4 activity and computations are inherited from the input regions (here, LGN) vs. being shaped by intrinsic, recurrent connectivity (see, e.g., [51]). To investigate this, in addition to regular (“Full”) simulations, we performed a number of simulations where recurrent and background connections were removed, so that neurons received LGN inputs only (“LGN only”). Using *in silico* voltage clamp (see Methods), we measured (**Fig. 4a**) synaptic excitatory currents at cells’ somata – from the LGN only, *I*_*LGN*_, and the total, *I*_*tot*_ – and found that, for preferred directions of 2 Hz drifting gratings, the fraction of *I*_*tot*_ contributed by *I*_*LGN*_ was 0.41+/−0.05 for excitatory cells, in good agreement with experiment (0.36+/−0.02 [17]). Also in agreement with experiment [17], the mean *I*_*LGN*_ was untuned, since individual LGN filters were mostly untuned [17] (but see [19, 20]), whereas mean *I*_*tot*_ and *I*_*sub*_ = *I*_*tot*_ — *I*_*LGN*_ (i.e., the current due to recurrent connections) were well tuned to the grating orientation (**Fig. 4b**), and F1 components (i.e., the amplitude of the mode at the stimulus frequency; see Methods) of all of these currents were tuned. Besides these features that were consistent with experimental recordings [17], one distinction was observed: the F1 component of *I*_*sub*_ was substantially smaller than that of *I*_*LGN*_ or *I*_*tot*_ (**Fig. 4b**) and the temporal dynamics of *I*_*sub*_ was in antiphase with that of *I*_*LGN*_ (**Fig. S6a**). This turned out to be a space clamp artefact, where a stronger *I*_*LGN*_ current increases the membrane voltage at the synapses, resulting in weaker driving force for recurrent excitation and, therefore, antiphase relationship of *I*_*LGN*_ and *I*_*sub*_; simulations where LGN current was removed from the recorded cells exhibited *I*_*sub*_ with no oscillations at the grating frequency (**Fig. S6a**). Thus, the overall picture was that the *I*_*LGN*_ mean was untuned whereas its F1 component was; the recurrent connections added current that was tuned overall, but was not time-modulated (whereas in the experiment [17] it is time-modulated); and the resulting total current was tuned in both mean and F1. The inhibitory currents were mostly untuned at the level of both the mean and F1 (**Fig. 4b**); the F1 did show a preference to orientation, but its magnitude was only a few percent of the mean and the current was not strongly modulated in time (**Fig. S6a**).

**Figure 4.**
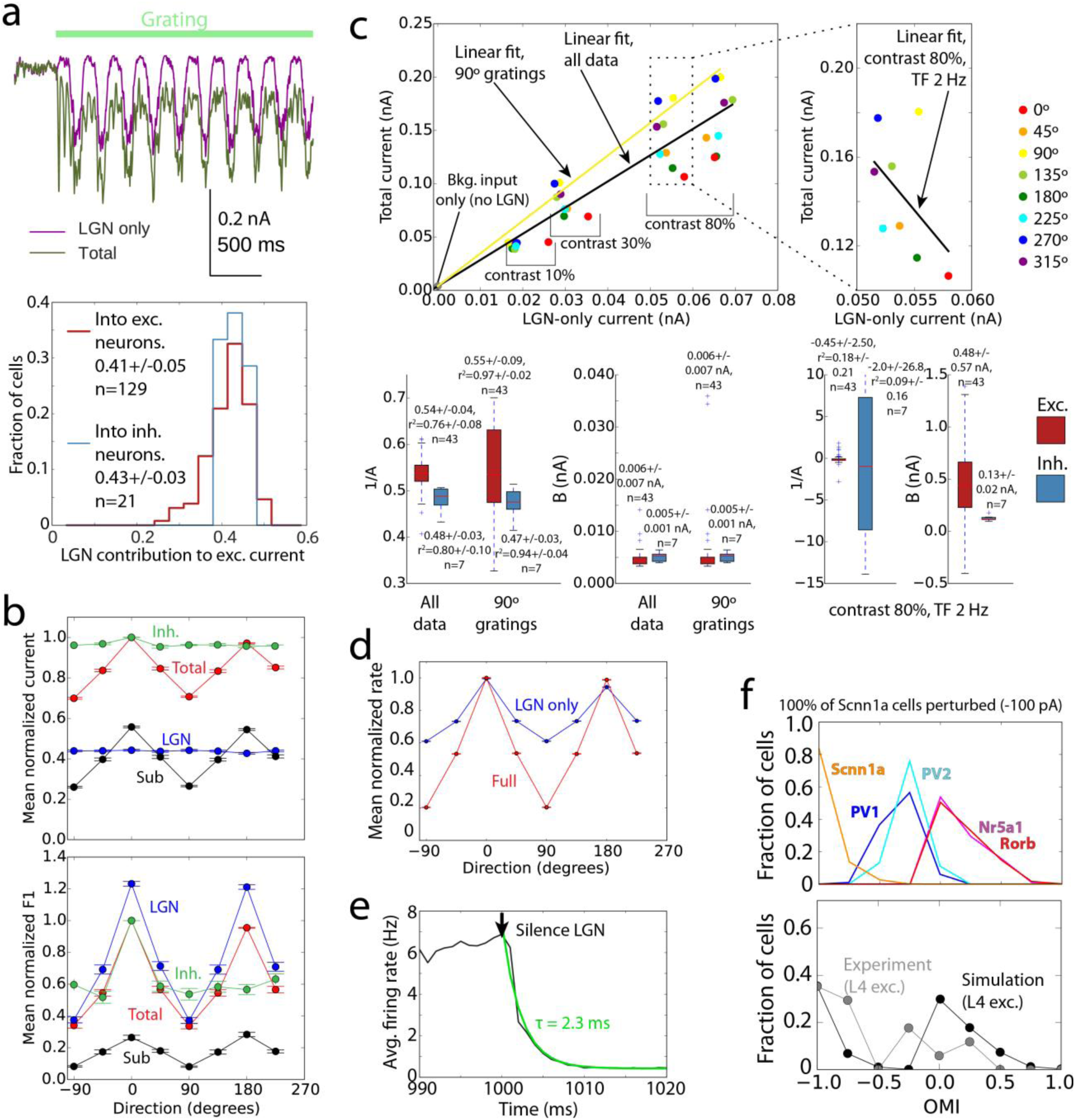
Mechanistic characterization of the model. (a) Cortical amplification of the LGN inputs. The excitatory currents (from the LGN only, as well as total) in biophysical cells were measured using voltage clamp recordings. Top – an example; bottom – distributions of LGN contribution to the total excitatory current across excitatory and inhibitory cells (computed for each cell as the average current over time and over all trials of the preferred orientation). (b) Tuning curves for the mean and F1 component of the total and LGN-only currents, and their difference (“Sub”, i.e., the cortical component), as well as inhibitory current. The data for each cell were normalized to the peak value of the “Total” and shifted so that the preferred direction is at 0 degrees; averages and s.e.m. over all recorded excitatory cells are shown (TF = 2 Hz, contrast 80%). The inhibitory currents were normalized and aligned to their own peak values, since their magnitude is significantly higher than that of excitatory currents. (c) Amplification of excitatory current. Top, the total current vs. the LGN-only current, for an individual Rorb cell (each point is an average over time and over 10 trials). Linear fits (*I*_*tot*_ = *A I*_*LGN*_ + *B*) are shown for data aggregated from all grating directions, TFs, and contrasts (black), for one selected direction (yellow), and for a fixed contrast and TF (i.e., representing a sample direction tuning curve; right plot). Bottom, summary of linear fits across all cells analyzed. (d) Tuning curves for mean firing rate in full network simulations (“Full”, red) and in simulations where all connections except the feedforward connections from the LGN were removed (“LGN only”, blue). The data for each cell were normalized to the peak value of the “Full” and shifted so that the preferred direction is at 0 degrees; averages and s.e.m. over all excitatory cells are shown (TF = 2 Hz, contrast 80%). (e) Simulations of responses to a drifting grating, with the LGN activity switched off at 1000 ms. The black curve is the firing rate averaged over all cells, models, and trials; green is the exponential fit. (f) Distribution of the optogenetic modulation index (OMI) by cell type in responses to gratings, for simulations of optogenetic silencing of the Scnn1a population (top). Combined distribution for all biophysical excitatory cells is compared to the experimental result (bottom).

Comparing mean *I*_*tot*_ and *I*_*LGN*_ of individual cells (**Fig. 4c**) for all grating conditions (i.e., not only the preferred one shown in **Fig. 4a**) and different contrasts, we observed a complex dependence. The relationship of *I*_*tot*_ with *I*_*LGN*_ was essentially linear along the contrast dimension (the quality of the linear fit,

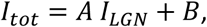

was r^2^=76% for excitatory cells for all conditions, and 97% for a specific grating direction), consistent with the prediction of an earlier, much simpler self-consistent model [51], but highly non-linear for the orientation dimension (r^2^=18%). The latter was the consequence of the mean of *I*_*LGN*_ being untuned to orientation, whereas the tuned current from recurrent connections made the mean of *I*_*tot*_ highly tuned (**Fig. 4b**). Due to the same reason, the amplification factor *A* was approximately 2 across all conditions (since *B* was very small on average, *A* can be thought of as the inverse of the LGN contribution described above; across all conditions *1*/*A* = 0.54+/−0.04, **Fig. 4c**), whereas for the preferred orientation it was 1/0.41=2.44 (as the LGN contribution at the preferred orientation was 0.41+/−0.05, **Fig. 4a**).

How do these relationships between currents shape the properties of the spiking output? We found that the OSI in L4 at the level of spikes was inherited from the combined LGN inputs to each cell [17], producing weakly selective output, and was strongly increased by recurrent connections (**Figs. 4d**, **S6b**). Generally, the relationship between the rates in Full and LGN only simulations (**Fig. S6c**) was not close to linear (attempting a linear fit

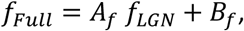

where *f*_*Full*_ and *f*_*LGN*_ are the firing rates in Full and LGN only simulations, and the parameters *A*_*f*_ and *B*_*f*_ are marked with the subscript *“f”* to distinguish them from the parameters of linear fit of currents above, we found r^2^=50% for all conditions and 65% for one grating direction for excitatory cells). The firing rates at fixed contrast and TF exhibited a much more linear relationship (r^2^=88%), but only locally – the good linear fit required negative values of parameter *B*_*f*_ (−4+/−3 Hz on average), which is non-physiological given that *B*_*f*_ is the firing rate of the cell under the conditions when the firing rate induced by the LGN-only input is zero. Thus, overall the relationship was non-linear, but close to linear past the firing threshold for LGN-only inputs. Interestingly, the overall effect of recurrent connections on the spiking output in our model was that of suppression, as the Full-network firing rates at non-preferred orientations tended to be smaller than LGN-only rates (**Fig. 4d** and **Fig. S6c**, top right), and only at the preferred orientations Full-network firing reached the same rates as in the LGN-only case. Consequently, *R*_*max*_ values were approximately the same in the Full and LGN-only cases (**Fig. S6b**).

To investigate the L4 circuit mechanisms further, we performed *in silico* optogenetic silencing of LGN inputs and of a subset of the circuit (see Methods). Despite the significant recurrent amplification, L4 activity shut down rapidly when the LGN spiking was silenced (**Fig. 4e**), in agreement with an analogous experiment [18]. Although the time course of activity decay was faster in the model (2.3 vs. 9+/−2 ms, likely due to the absence of excitatory stimulation from other cortical regions), otherwise the effect of LGN silencing was the same. The widespread and powerful intracortical inhibition appears to be the most plausible driving force behind this effect. We then separately silenced the Scnn1a population of excitatory cells in simulations and conducted analogous experiments *in vivo* (using Archaeorhodopsin-mediated silencing on randomly selected trials during presentation of TF = 2 Hz drifting gratings, while extracellular multielectrode recordings were performed, see Methods). Results were characterized by converting firing rates with (*f*_*p*_) and without (*f*_*control*_) perturbation to the optogenetic modulation index (OMI),

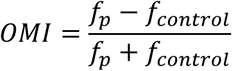

Simulations qualitatively agreed with experiment (**Figs. 4f, S7a, b**) in terms of the OMI distribution over L4 excitatory cells (few inhibitory cells were recorded experimentally), which was approximately bimodal, with one “lobe” concentrated close to −1 (totally silenced neurons) and the other near 0 (weak or no effect). Due to challenges of recording form the thin layer 4, a small number of cells was obtained in the experiment, but they all showed a consistent trend of OMI values belonging to one of these two lobes (**Fig. S7a**). Whereas weak and/or sparse silencing of Scnn1a cells in simulations did not result in a two-lobe distribution, a strong and dense silencing did (**Fig. S7c**), consistent with the experimental perturbation that attempted to silence as many Scnn1a cells as possible. Simulations showed (**Fig. 4f**) that the left lobe in the OMI distribution was due to near-complete silencing of Scnn1a cells. This silencing reduced the amount of excitation in the network and resulted in moderate decrease of firing of inhibitory cells. The net effect is that the firing rates of the other excitatory cell types (Rorb and Nr5a1) remained almost unaffected – yielding the OMI lobe with values close to zero.

Finally, we investigated the effect of the logics of connections between L4 excitatory cells on the neural activity and computation. In our regular models, a “like-to-like” (“L”) rule [25, 26, 24, 29] was used for both the connectivity and synaptic weights of excitatory-to-excitatory connections, based on the anticipated orientation tuning of the cells (**Fig. 1c-e**); because the rule applied to both synaptic connectivity and amplitude distributions, we refer to this set as “LL”. We then studied three alternative sets of models. In one, both the connectivity and weights were randomly (“R”) assigned independently of cell tuning (“RR”). The remaining two sets had the random rule applied to connectivity and like-to-like to weights (“RL”), or vice versa (“LR”). Each set consisted of 3 models. Besides the “R” or “L” rules, everything else was exactly the same between the four sets, including the probabilistic distance-dependent connectivity (**Fig. 1c**).

Synaptic weights for all models were tuned following the same procedure (tuning only involved scaling of weights uniformly across population, thus not affecting the “L” vs. “R” property, which applies to individual cell pairs), resulting in similar levels of activity for training stimulus (grating) in all models. We then assessed how functional properties differed. We found that OSIs were extremely reduced in the RR set, with LR being only slightly better tuned, whereas RL came close to the original well-tuned LL set (**Fig. 5a**). Because the mean LGN current was untuned in all models, the critical amplification of orientation selectivity (**Fig. 4b-d**) came from the lateral L4 connections, which were well tuned in LL and RL sets and poorly tuned in LR and RR sets (**Fig. 5b**). As a result, the total current as well as spiking output was well tuned in LL and RL sets and barely tuned in LR and RR **(Fig. 5)**. This suggests a bigger role of synaptic weights than connection probability in shaping network responses, which can be easily understood since the “L” rule for weights results in low contributions from non-like-to-like connections, even if such connections are present, thus effectively enforcing the “L” type connectivity.

**Figure 5.**
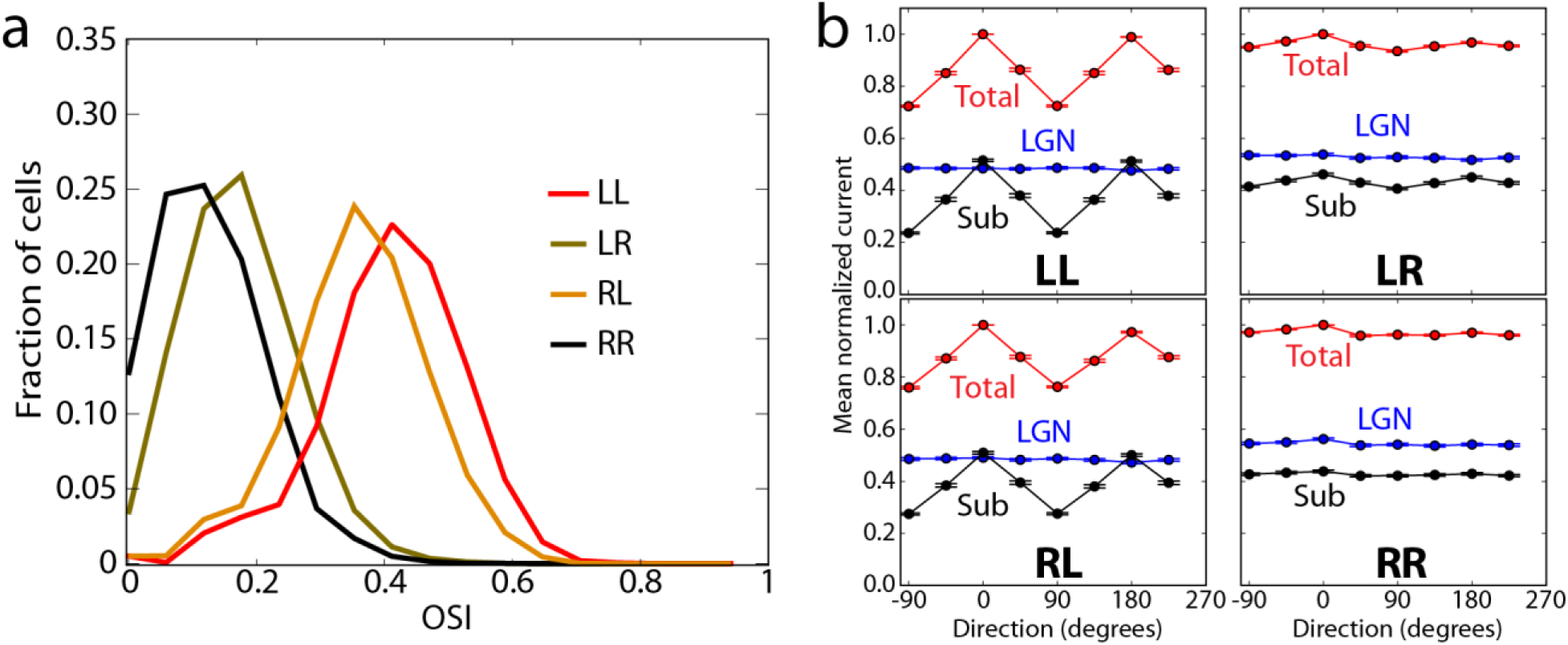
Like-to-like vs. random connectivity and synaptic weights. (a) Distribution of OSI for biophysical excitatory cells for the LL, RL, LR, and RR cases. (b) Tuning curves for the mean total and LGN-only currents, and their difference (“Sub”, i.e., the cortical component). The data for each cell were normalized to the peak value of the “Total” and shifted so that the preferred direction is at 0 degrees; averages and s.e.m. over all recorded excitatory cells are shown (TF = 2 Hz, contrast 80%).

### Performance of a simplified model

How does the ability to reproduce experimental recordings depend on the level of granularity of the model? To address this question, we built two versions of a radically simplified model, where biophysical neurons and bi-exponential synapses were replaced by LIF neurons and either instantaneous “charge-dump” synapses (using NEURON’s IntFire1 function) or synapses with exponential time dependence of synaptic current (NEURON’s IntFire4 function). These all-LIF models used exactly the same cell-to-cell connections and inputs as the biophysical models and were optimized using the same protocol (see Methods).

The results of the all-LIF simulations were qualitatively similar to the results of biophysical simulations (**Fig. 6**), but specific quantitative distinctions were obvious. Although the overall appearance of responses (**Fig. 6a**) and mean values of spontaneous rates and *R*_*max*_ were similar to the biophysical case, the OSI values were elevated in the IntFire1 case (**Fig. 6b**). Most noticeable, the gamma oscillation was largely absent in the IntFire1 models (**Fig. 6c**). This was likely due to instantaneous synapses in IntFire1, since gamma oscillations [11, 12, 13, 14, 15] are thought to be strongly dependent on the properties of inhibitory perisomatic synapses [11, 14], and especially their time constants. Thus, removal of the appropriate synaptic kinetics from the large and distributed network model like ours may be expected to disrupt gamma oscillations. By contrast, the IntFire4 model employed a simple form of synaptic kinetics and produced oscillations in the range of 10-20 Hz, i.e., close to ∼20 Hz gamma oscillation in the biophysical model (**Fig. 6c**).

**Figure 6.**
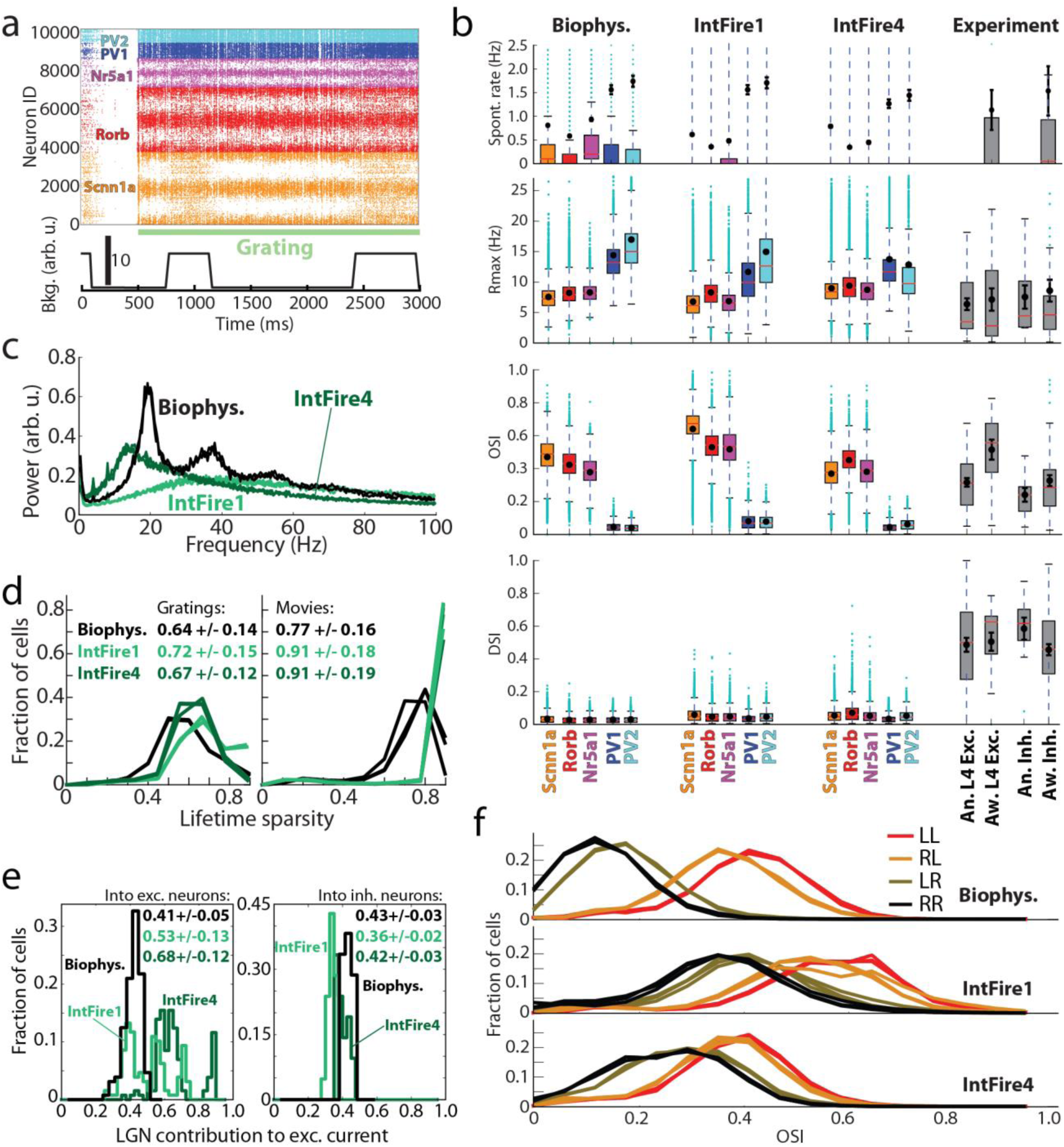
Comparison of biophysical and all-LIF simulations employing the IntFirel or IntFire4 models. (a) An example spike raster in response to a drifting grating in an all-LIF IntFire1 simulation. (b) Spontaneous rate, *R*_*max*_, OSI and DSI by cell type. (c) Spectra of multi-unit activity (weighted by *1*/*r,* where *r* is the distance from the cell to the center of the system; this is used as a proxy to LFP). (d) Distributions of lifetime sparsity of responses to gratings and movies, averaged over 10 trials. The data are for three directions of a grating (0, 45, and 90 degrees) and for three movies. (e) Distributions of LGN contribution to the total excitatory synaptic inputs across excitatory and inhibitory cells (in the IntFire1 and IntFire4 cases, this is computed for each cell as the average synaptic input over time and over all trials of the preferred orientation; see Methods), for TF = 2 Hz drifting gratings. (f) Distribution of OSI for excitatory cells for the LL, RL, LR, and RR cases.

**Figure 7.**
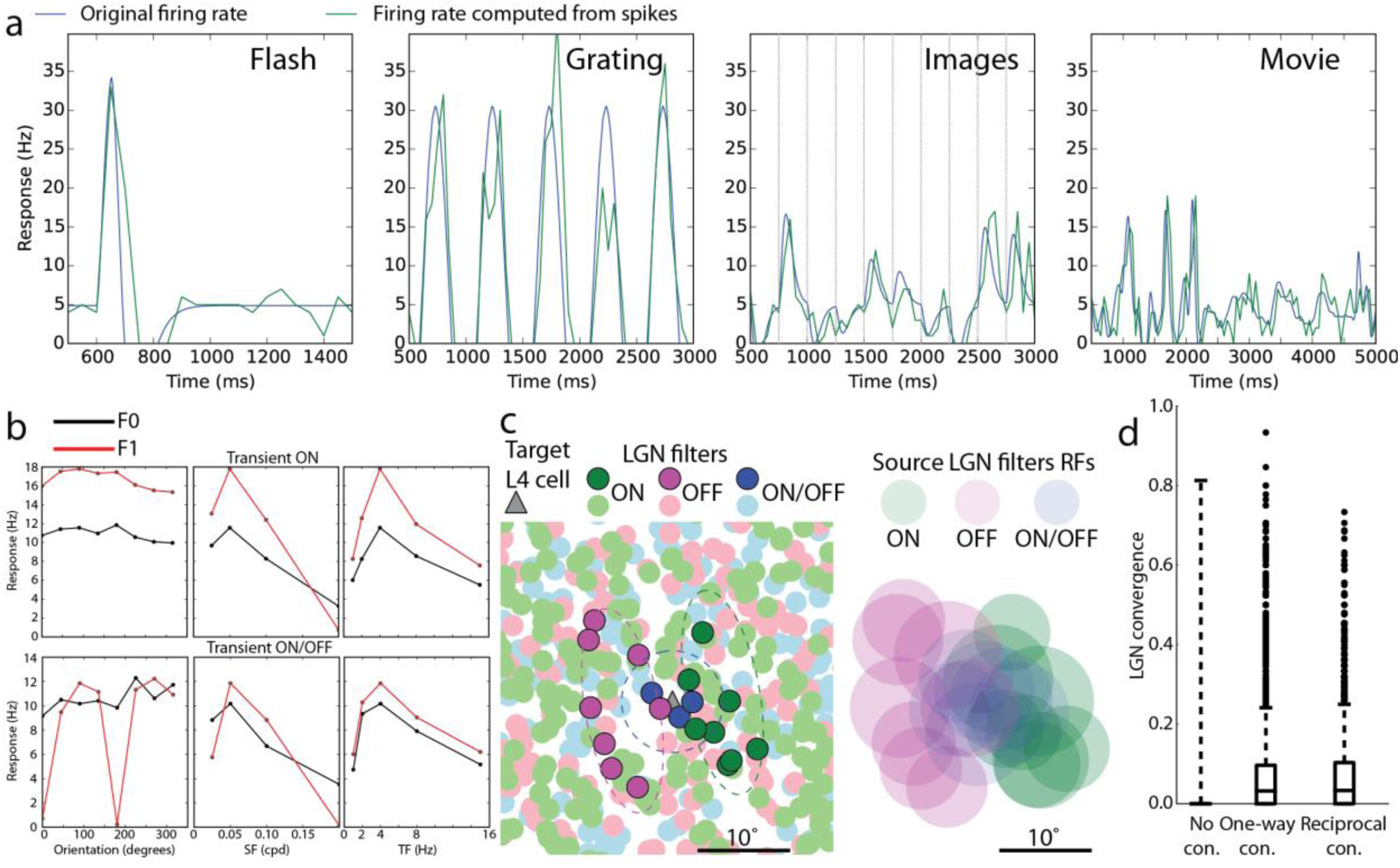
LGN filters. (a) Example responses of a single filter to visual stimuli, as a time-dependent firing rate that is the filter output (blue) and the firing rate computed from generated spike trains, averaged over all trials (green). For clarity, the first 500 ms of each trace are not shown; during this interval, the rate is set to a constant corresponding to the level of spontaneous activity of the filter. The panel for images contains responses to 10 images shown in a sequence, 250 ms each. (b) F0 and F1 components of the responses to gratings for two example LGN filters. Tuning curves to orientation, SF, and TF are shown. The data points are averages from generated spike trains over time and over trials. (c) Connecting LGN filters to L4 cells. Geometry in the visual space is illustrated. The left panel shows centers of all filters present in a portion of the visual space around the mapped position of an example excitatory cell from L4. The dashed lines correspond to the “lasso” subfields around one illustrative L4 cell, used to capture input LGN filters of the ON, OFF, and ON/OFF type. The filters that are selected to send inputs to this L4 cells are in deep color; all other filters are dimmed. On the right, the same L4 cell with the filters selected to provide inputs to it are shown. For the filters, the approximate size of their receptive subfields is illustrated (a single subfield for ON or OFF filters and two subfields for ON/OFF filters; the radius of each RF circle is 2*σ*_*c*_). (d) Convergence of LGN connectivity onto L4 cells that are not connected to each other (“No con.”), one-way connected (“One-way con.”), and reciprocally connected (“Reciprocal con.”). For each pair of L4 cells, the LGN convergence is defined as the number of LGN filters that connect to both cells divided by the sum of the numbers of LGN filters connected to each of the cells. The data are aggregated from three L4 models.

Further distinctions were observed for sparsity values (**Fig. 6d**). The overall trend of higher sparsity for movies than for gratings was reproduced by both all-LIF models, but they both also exhibited higher sparsity values than the biophysical case and the experiment (**Fig. 3f**). Perhaps the most significant difference with the biophysical model was observed for the magnitude of cortical amplification (**Fig. 6e**) for excitatory cells (interestingly, inhibitory cells did not show substantial differences). The LGN contribution was 0.41+/−0.05 in the biophysical model and 0.36+/−0.02 in experiment [17], whereas it was 0.53+/−0.13 for IntFire1 and even higher, 0.68+/−0.12, for the otherwise more realistic IntFire4. The distributions of amplification values were also quite different in all-LIF models (**Fig. 6e**), exhibiting multiple peaks – apparently for the multiple excitatory cell types – instead of a single peak in the biophysical model. Finally, the functional connectivity had the same consequences in the all-LIF models as in the biophysical case – the orientation tuning was high for models with the LL or RL rules, and low for LR and RR (**Fig. 6f**). On the other hand, the small difference between the LL and RL cases, observed in the biophysical model, was mostly eliminated, and overall the OSIs were higher for the IntFire1 model and, in the RR and LR cases, for IntFire4 as well.

## DISCUSSION

The promise of data-driven neuroscience modeling is in harnessing computing power to establish a platform for discovery that would work hand-in-hand with experiments. For that promise to materialize, models need to be biologically realistic in recapitulating knowledge about brain structure as well as in reproducing *in vivo* activity and computation. Doing either is difficult, and doing both together is more difficult yet, and has not been widely practiced (though, see, e.g., [2, 3, 4]). We described here a model of the L4 in mouse V1 that was built using a high degree of biological realism and simulated in a framework of an *in silico* visual physiology experiment. We asked how well the model reproduces activity observed *in vivo* across a variety of stimuli, which mechanisms underlie the activity and computation in the modeled L4 circuit, and how the levels of simplification used in the model affect its performance.

Despite the small training set of stimuli for which our model was optimized – a gray screen and a single grating presentation for 0.5 s – it generalized well to a large test set of stimuli. The model reproduced major features of *in vivo* observations with respect to, e.g., the magnitude of responses to gratings, orientation selectivity, prominence of gamma oscillations, long-tailed distributions of firing rates, lifetime sparsity for gratings and movies, magnitude of cortical amplification, and effect of optogenetic perturbations of the LGN or the Scnn1a population in L4 (**Figs. 3, 4**). Such an agreement across stimulus classes and types of observation is remarkable given that, although the model included a significant degree of biological realism, many simplifications were used: neurons possessed active conductances only at the soma, the variety of neuron models was limited to five unique single cell morphologies and sets of membrane conductances, synapses had no short-term plasticity, LGN inputs were simplified, connectivity was established via simple probabilistic rules, most known interneuron cell types [33] were absent, and in fact the influence of most of V1 (L1, L2/3, L5, and L6) and the rest of the brain was reduced to extremely simple background states. This may suggest that many of L4 computations are produced by its local network, and other layers may play a primarily modulatory role – such as gain control exerted by L6 [52] – as far as L4 activity is concerned.

Perhaps even more instructive than successes, some deficiencies were observed too. In terms of reproducing *in vivo* activity, the most important issues were the absence of direction selectivity and too fast responses to full-field flashes (**Fig. 3b,e**). Both appear to be due to the extreme simplicity of the LGN inputs – in particular, the absence of sustained LGN responses in the model. This provides an immediate direction for improving the model, especially based on the more recent experimental results [23], which is the aim of ongoing work on the next model generation.

In applying the model to study mechanisms underlying the operation of the L4 circuit (**Figs.4, 5**), we found the following principles. Tuning in L4 cells arose from combination of convergent LGN inputs, but to reach physiological levels of selectivity, the functional like-to-like connectivity was essential (**Figs. 4d, S6b, 5**). The effect was especially sensitive to synaptic weights, and less so to connectivity alone (**Fig. 5**), because like-to-like weights effectively enforce like-to-like connectivity by scaling down contributions from non-like-to-like connections. These results suggest that like-to-like rules, experimentally observed in L2/3 [25, 26, 24, 29], are likely to be found in L4 as well, and may play an essential role in determining functional information processing in the cortex.

The amplification of excitatory current in excitatory cells due to recurrent connections was by 2-3-fold (**Fig. 4a, b, c**), in quantitative agreement with experimental measurements [17]. The amplification was linear [51] along the contrast dimension (possibly, in part due to our LGN filters being linear), but highly non-linear along the orientation dimension (**Fig. 4c**), since the total current was tuned to orientation, whereas the LGN current was untuned [17] (**Fig. 4b**). Thus, overall the network amplification was extensive and non-linear both at the level of currents and at the level of spiking output (**Fig. S6c**). Despite such strong excitatory amplification, the whole circuit was controlled by even stronger inhibition, which readily shut down the activity in the absence of the LGN input [18] (**Fig. 4e**). Optogenetic suppression of a subpopulation of excitatory L4 neurons (Scnn1a) resulted in a clearly bimodal distribution of firing rate changes both in simulation and in experiment (**Fig. 4f**), demonstrating a compensatory effect in the circuit, where only the directly targeted Scnn1a cells were shut down, whereas other cells approximately maintained their activity levels.

In terms of the mechanistic characterization of the circuit, another important insight again follows from the model deficiency: we observed that the cortical component of the excitatory current (“Sub”) into excitatory neurons was tuned to orientation, but poorly modulated in time (**Figs. 4b, S6a**), whereas in the experiment it was strongly modulated at the grating frequency and closely matched the phase of the LGN input [17]. In the model, this “Sub” component is tuned because of the like-to-like connectivity rule, where cells receiving similarly oriented LGN inputs have higher connection probability and stronger synapses. However, this rule does not take phase into account, and therefore, for a given target cell, all source cells have different phases, explaining why the “Sub” current was not modulated at the grating frequency. The fact that in the experiment “Sub” current is modulated and is in phase with the LGN current indicates that the like-to-like connectivity is more sophisticated and includes also the phase information [17]. This is consistent with reports in the literature [25, 24] that similarity in orientation preference is a good but not the best predictor of connectivity and with theoretical ideas (e.g., [28]) that lateral connections in the cortex are optimized to enhance features, such as extended lines in V1.

A further simplification of our network model, replacing all biophysical neurons by LIF units, preserved most general trends in features of neuronal activity, but the quantitative agreement with experiment suffered. The levels of orientation selectivity and sparsity of responses were altered (**Fig. 6b, d, f**). The oscillations at the level of population activity were eliminated when instantaneous synaptic kinetics was used and partially rescued with non-instantaneous synapses (**Fig. 6c**). Although further exploration is necessary to find out how one can match the oscillations spectrum of the biophysical model with a simpler network model, this result supports the idea that synaptic kinetics is important for generating oscillations in the relevant regime (e.g., [11, 14]). Perhaps the most substantial difference with the biophysical model was that the cortical amplification was not captured well by all-LIF models (**Fig. 6e**), hinting that mechanisms shaping activity and computation in the L4 circuit may not be well represented by these simpler models. Clearly, many modifications of the simpler models are possible that may improve performance, and our results do not rule out that point-neuron simulations can be adequate for studying cortical activity and function – in fact, our results are promising since many of the trends of activity were reproduced qualitatively. But, they do suggest that careful characterization of the level of representation used for modeling may be important and that biological detail may matter for faithful representation of a variety of observables.

Overall, results reported here are encouraging in that, despite significant simplifications, many aspects of activity and computation in the L4 circuit were captured by a data-driven model constructed in a bottom-up fashion. We suggest that systematic data-driven modeling oriented towards mimicking *in vivo* physiology experiments will be an important approach to take advantage of the modern large-scale data collection efforts. Our software code, the model, and simulation results are made publicly available (see SI) to enable further efforts in modeling *in vivo* activity and function.

## DATA AVAILABILITY

Data generated or analyzed during this study are included in this published article in its supplementary files; as specified in the SI, larger volumes of the data are made available online, and additional raw data is available on request (per cost for data medium and shipping). Modeling and analysis also involved data that are already publicly available (Allen Brain Observatory [53] and Allen Cell Types Database [31]).

## CODE AVAILABILITY

The software code used to build models, perform simulations, and perform analysis is included in this published article in its supplementary files.

## AUTHOR CONTRIBUTIONS

A.A. and C.K. designed the overall study. J.B., N.dC., S.dV., D.D., S.D., N.W.G., R.I., T.J., J.L., B.L., S.M., S.R.O., R.C.R., G.S.-L., S.A.S., M.S., Q.W., and J.W. contributed to the design of models and simulations, supplied data, and provided crucial advice. A.A., Y.N.B., N.C., D.F., N.W.G., S.G., R.I., Z.W., and Z.X. developed modeling software, built models, and carried out simulations. N.dC., D.D., S.D., L.L., and S.R.O. performed experiments. A.A., Y.N.B., M.B., N.dC., D.D., S.D., N.W.G., S.G., R.I., G.K.O., S.R.O., Z.W., and Z.X. analyzed the data. A.A. and C.K. wrote the manuscript with comments from all authors.

## ACKNOWLEDGEMENTS

We are grateful to Gabe J. Murphy for helpful discussions. We thank the Allen Institute founders, Paul G. Allen and Jody Allen, for their vision, encouragement and support.

## METHODS

### Model building and simulation details

All simulations were performed with the parallelized code (available in *Supplementary Files)* written in python 2.7 and employing NEURON 7.4 [34] as a simulation engine. Simulations were carried out typically on 120 CPU cores (Intel Xeon CPU E5-2620 v2, 2.10GHz; 128 GB RAM per a 24-core node), requiring ∼1 hour to simulate 1 s.

#### Network Composition

The network model was constructed from five biophysically detailed models of individual neurons [30] from an early version of the Allen Cell Types Database [31], and two models of leaky-integrate-and-fire (LIF) neurons (one for excitatory type and one for inhibitory type). The biophysical models employed passive conductances in the dendrites and 10 active conductances in the soma, including Na^+^, K^+^, and Ca^2+^ conductances, as described in detail in [30]. The five models represented three types of excitatory L4 neurons (expressing the genes Scnn1a, Rorb, and Nr5a1) and two types of fast-spiking parvalbumin-expressing (PV) interneurons. Cells expressing these markers comprise the majority of neurons in L4 of V1. These models consisted of 264 (Scnn1a), 141 (Rorb), 101 (Nr5a1), 121 (PV1), and 91 (PV2) compartments. The LIF neuron models were implemented using the IntFire1() function of NEURON, which contains two parameters – the time constant and the refractory period. The former was set to the average time constant from the corresponding biophysical models (from Scnn1a, Rorb, and Nr5a1 for the excitatory LIF neuron and PV1 and PV2 for the inhibitory LIF neuron); the latter was 3 ms for both LIF models. All the models are available in the *Supplementary Files* (see SI).

The 45,000 cells in the network model were distributed as follows: 3700 for Scnn1a, 3300 for Rorb, 1500 for Nr5a1, 800 for PV1, 700 for PV2, 29750 for excitatory LIF, and 5250 for inhibitory LIF. The cells were distributed in a cylinder 100 μm in height; the biophysical cells occupied the inner core with the 400 μm radius, and the LIF cells, the outer shell with the radii from 400 μm to 845 μm.

Three independent models were generated using different random seeds. These three models were used for each connectivity case (LL, LR, RL, and RR, see Main Text). For each model, the recurrent connectivity and synaptic weights (see below) differed across the LL, LR, RL, and RR cases, since different connectivity/weight rules were applied; everything else was identical across these four cases.

#### Connectivity within the Network

For the purposes of establishing recurrent connections, all neurons were considered to belong to two groups – excitatory (E; Scnn1a, Rorb, Nr5a1, and excitatory LIF) and inhibitory (I; PV1, PV2, and inhibitory LIF). Four types of connections were established: E-to-E, E-to-I, I-to-E, and I-to-I. The probability of connection was chosen as a product of a function dependent on the distance between the somata of the two cells and the function dependent on the difference of the assigned preferred orientation angle (see **Fig. 1**). The latter function was constant, 1.0, for E-to-I, I-to-E, and I-to-I, whereas, for E-to-E it differed depending on the model we assumed. The models with like-to-like connection probability (LL and LR, see Main text) employed a linear function, starting at 1 for the zero difference in the preferred orientation angle, and equal to 0.5 at 90 degrees difference; for the models with random connection probability (RL and RR), the function was set to one. The distance dependent function was linear, with the peak at zero distance (0.34 for E-to-E and 0.26 for E-to-I in the case of LL and LR models; 0.255 for both E-to-E and E-to-I in the case of RL and RR models; and 1.0 for I-to-E and I-to-I in all cases). The linear decay was determined by the distance at which the function became zero (300 μm for E-to-E and E-to-I, and 160 μm for I-to-E and I-to-I). For established connections, the number of synapses was selected randomly with uniform probability between 3 and 7.

The numbers of synapses from different sources per neuron are not well established for the mouse visual cortex, but are known approximately for cat [40], where, for the L4 excitatory cells (spiny stellate cells projecting to L2/3 or L4, and for pyramidal cells), the number of incoming synapses is ∼5,000-7,000. Out of those, e.g., for pyramidal cells, ∼1,100 synapses come from L4 excitatory cells, ∼2,550 from excitatory cells in other layers, and ∼2,000 are excitatory with unassigned source. Assuming the same distribution of sources for unassigned synapses, one can expect that ∼1,700 excitatory synapses are from L4 sources. Note that the number of synapses from LGN is ∼100-150 in the cat cortex. Converting these numbers to fractions, one finds that ∼25-30% of all synapses are from the L4 excitatory sources, 55-60% from other excitatory sources, and 15-20% from inhibitory sources. For the inhibitory cells in L4, the fractions are similar, but the total number of synapses is smaller (∼3,250 synapses for basket cells).

We assumed similar fractions for the mouse visual cortex, except the number of synapses from the LGN, for which we experimentally found a much larger number, ∼1,000 per cell (see Main Text). Assuming somewhat smaller number of synapses for mouse cells than for cat cells, we posited (**Table 1**) for the purposes of our model that pyramidal cells in L4 of V1 receive ∼6,000 synapses total, with ∼1,000 (17%) coming from LGN, ∼2,000 (∼33%) from L4 excitatory sources, and ∼400 (∼7%) from L4 PV^+^ inhibitory sources. The rest was not modeled explicitly (except for the background, see Main Text and below), as it comes from other L4 inhibitory sources (PV^−^ interneurons – presumably, another 7%, i.e., ∼400 synapses) and from excitatory sources from other layers and brain regions (∼2,200, i.e., 36%), both of which were not represented in the model. For the synapses from the LGN, we did not model explicitly all the LGN cell types (see below), and thus restricted the number of LGN synapses to ∼600 out of the expected 1,000 total. For simplicity, we assumed numbers for synapses from LGN and L4 neurons to the PV+ cells to be similar in our model to the numbers used for pyramidal cells. Due to the randomness of connections and of the number of synapses per connection, the actual number of synapses from L4 sources for each cell varied within ∼10% from the average (**Fig. S1a**).

**Table 1.** Assumed average numbers of synapses per an excitatory cell in L4 of mouse V1. See text for details.

| From LGN | From L4 excitatory cells | From L4 PV<sup>+</sup> interneurons | From L4 PV<sup>-</sup> interneurons | From neurons in other layers and brain regions | Total |
| --- | --- | --- | --- | --- | --- |
| 1,000 | 2,000 | 400 | 400 | 2,200 | 6,000 |

Thus, each cell in our simulations received on average ∼3,000 synapses from other excitatory and inhibitory cells in the L4 model and LGN sources. After building our models, including connectivity, we found that, for recurrent connections among the L4 cells, excitatory neurons received connections from 490 +/−20 and sent connections to 480 +/−20 neurons on average, whereas inhibitory neurons received connections from 490 +/−20 and sent connections to 530 +/−20 neurons on average. In addition, each cell also received a small number of “background” synapses, as described below.

Synapses were distributed on the dendritic trees of the target cells randomly within certain distance constraints, on the soma, basal, or apical processes, according to the literature (e.g., [32, 33, 40, 54, 55, 56, 57, 58]). Namely, all synapses to PV1 and PV2 cells were distributed on the soma and basal dendrites without limitations. For Scnn1a, Rorb, and Nr5a1 cells, excitatory synapses from L4 and background were placed on basal and apical dendrites, 30 to 150 μm from the soma, excitatory synapses from LGN were placed on basal and apical dendrites, 0 to 150 μ from the soma, and inhibitory synapses were placed on the soma and basal and apical dendrites, 0 to 50 μ from the soma.

#### Model of background inputs

Whereas the recurrent connections within L4 and the LGN inputs (see below) were modeled explicitly, the connections from the rest of the brain were represented using a simple model of “background” traveling waves (see Main Text). For producing the waves, we distributed 3,000 Poisson spike generators in the x-y space covered by the model (the L4 plane), and drew random connections from these generators to the modeled L4 cells if the distance between a generator and a cell’s soma was within 150 μm, allowing between 18 and 24 such connections per cell. This approximation of a small number of connections (much fewer than what is expected for incoming connections per L4 pyramidal cell from outside L4 or LGN) was chosen for simplicity and for reducing computing expense; in the absence of heterogeneity in the population of sources, providing a much larger number of connections does not appear necessary.

The background sources produced Poisson spike trains from a time dependent firing rate, *f(x, y, t),* which was controlled by the plane traveling waves (see **Fig. 2**). The waves were pre-generated from a different random seed for each simulation trial. The direction of each wave’s movement in the x-y plane was randomly selected, its width in the direction of movement was kept constant in time, in the perpendicular direction the wave was infinite, and its magnitude was also constant and randomly selected to be between 5 and 15 Hz. In the absence of the wave, *f(x, y, t)* for each spike generator was set to zero, and it sharply rose to the value given by the wave’s height when the wave moved through the (x, y) location of the generator. The waves were produced so that they rarely overlapped, and thus the firing rate of the spike generators mostly exhibited periods of silence and periods of constant firing output, as waves swept through each generator’s location (see **Fig. 2**).

The utilization of background traveling waves furnished a simple and computationally efficient analogue of different cortical states. To approximate physiological observations with the choice of the wave parameters, we performed a small set of patch-clamp recordings in the pyramidal cells (n=3, targeting L2/3 neurons in V1 for easiness of access) of anesthetized mice. In the neural responses during spontaneous activity (gray screen; **Fig. S1b**), we observed clearly distinct intervals of rest and baseline depolarization, and found that on average the duration of depolarized states was 700+/−300 ms and the interval between the end of one such state and beginning of the next was 1,100+/−600 ms, overall consistent with observations illustrated in the literature (e.g., [43, 42]). Clearly, this is only one measurement under particular conditions, and, generally speaking, the actual characteristics of the cortical states, such as duration of the hyperpolarized state, interval between such states, and also potentially gradations in the degree of depolarization or hyperpolarization, would depend on the type of modulation one considers. These characteristics may exhibit a variety of time scales. Nevertheless, it appears that largely these cortical states may be modulated on the time scale of 1,000 ms or longer, by the order of magnitude. Therefore, for simplicity, for the L4 model we generated background waves that lasted from 200 ms to 1,200 ms (i.e., 700 ms on average) and were separated by intervals of 250 ms to 1,750 ms (i.e., 1,000 ms on average) – both parameters being drawn randomly for each wave from uniform distributions. The spatial extent of each wave was set to 2,000 μm, and the propagation speed was set based on that and on the randomly selected duration of the wave.

The remaining parameters were the number of synapses per connection from a spike generator to a L4 cell and the conductance amplitude of these synapses at the synapse site. The amplitudes were set to a single number for cells of one type; the variation in the resulting postsynaptic potentials/currents (PSPs/PSCs) was due to varying synapse number per connection and random placement of synapses on the dendritic tree (see below for a general description of the choice of synaptic weights). First, the weights for the background synapses were set so that the *V*_*m*_ of target cells was elevated by 10-15 mV from rest during the wave condition (which was the amount of elevation we observed between the depolarized and rest states in the patch clamp experiments), in purely feedforward regime in absence of any other connections. The weights were then further adjusted to reproduce the spontaneous firing rates in the context of the full network (see below).

The number of synapses per background connection was the remaining parameter that we used to generate a long-tailed distribution of background input strengths, which we found to be important for producing a skewed (log-normal-like) distribution of firing rates (see **Fig. 3a**), as observed experimentally [5]. Specifically, for each L4 cell, the numbers of synapses for every background connection to that cell were the same, but differed between cells. These numbers were drawn from a long-tailed distribution heavily weighted towards 1 synapse, but permitting up to 16 synapses (**Fig. S1c**). As a result, most L4 cells received similar amount of background excitation, but small fractions of cells received much higher amounts of background excitation.

#### Synaptic characteristics

Bi-exponential synapses (NEURON’s Exp2Syn) were used for biophysical and instantaneous synapses for LIF target cells; the reversal potential was −70 mV for inhibition and 0 mV for excitation. For simplicity, synapses did not employ short-term or other plasticity rules and had 100% release probability. Although presence of such properties is well documented for individual cortical synapses, especially *in vitro,* their role in functional properties *in vivo* is not well understood. Short-term plasticity may be less prominent *in vivo* than *in vitro* (due to saturation to baseline levels) [8], whereas non-deterministic nature of individual synapses is somewhat alleviated by the fact that multiple such synapses are typically present for each connection. The role of these properties *in vivo* is an important subject for future investigation. In the current study, neglecting these properties lead to significant conceptual simplifications as well as savings in computing power, and the generally good agreement of the neural activity in the simulations with the experiment (see Main Text) suggest that the role of these properties may be subtle and not straightforward to analyze. As a side observation, one should note that, because the modeled synapses were deterministic, the synaptic weights we used are probably underestimated in terms of the conductance values at the synapse location. However, the postsynaptic effects at the soma – peak PSPs and PSCs – were found to be consistent with the experiment (see below).

No complete resource exists yet with a characterization of synaptic weights and kinetics in L4 of mouse V1, and the literature on the subject of synaptic weights in general is large and somewhat disparate in the details of experimental methods used and features recorded and reported (such as post-synaptic potentials, or PSPs, vs. post-synaptic currents, or PSCs, definitions of time constants for PSP or PSC kinetics, etc.). Therefore, choices of synaptic weights in our models were guided only approximately by the existing literature on the synaptic properties of the cortex in various species (e.g., [25, 45, 47, 59, 60]), and for various cortical areas and layers (e.g., somatosensory [47] and anterior cingulate [45] cortices). Because of the lack of a consistent benchmark, we aimed at restricting the synaptic weights and time constants within approximately an order of magnitude of the values reported across a variety of experimental studies. The kinetic parameters were fixed in the models, whereas the weights were allowed to be varied globally (such as, e.g., uniformly scaling weights of all excitatory synapses on Scnn1a biophysical cells by a multiplicative constant, etc.) at the optimization stage. The optimization stage aimed at reproducing firing rates to a grating, spontaneous activity, and avoid epileptic-like time-locked activity of all cells (see Main text). Once the weights were scaled so that these aims were approximately reached, they were kept constant throughout all simulations for a given model.

The synaptic weights were given constant values for each connection type (namely, four source types – excitatory from L4, inhibitory from L4, excitatory from LGN, and excitatory from background – to the seven target types – Scnn1a, Rorb, Nr5a1, PV1, PV2, and excitatory and inhibitory LIF), which for the biophysical cells corresponded to a constant value of the peak conductance at the synapse site. For the models where like-to-like synaptic weights were employed (LL and RL, see Main Text), the uniform synaptic weights from above were further multiplied by a factor *F*_*w*_ that depended on the difference between the assigned preferred orientation angle between the two cells:

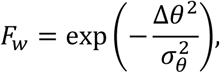

where Δ*θ* is the angle difference (defined within 90°), and *σ*_*θ*_ = 50° was used in all cases.

Despite weights at the synapse location being constant for most connection types, the actual PSPs or PSCs at the soma were widely distributed, because synapses were randomly distributed on the dendritic trees. The peak somatic PSCs for E-to-E synapses (**Fig. S1d**), for example, were observed in the model to be within 2 orders of magnitude, consistent with experimental findings [54]. Their distribution on the log scale was wide – i.e., similar to a skewed, lognormal distribution (or representable as a sum of a few lognormal distributions) described in the literature [5]. These distributions were very similar between the LL, RL, LR, and RR models, half of which employed constant E-to-E synaptic weights and other half used *F*_*w*_-modulated weights.

After models were optimized and simulated, we extracted a subset of synaptic connections (n=100, each containing 3 to 7 synapses, as explained above) from each model for each connection type (E-to-E, E-to-I, I-to-E, I-to-I). We computed the PSCs and PSPs in voltage clamp and current clamp, respectively, and analyzed their peaks and time course (**Fig. S2**). For the current clamp measurements, current was not injected, thus allowing for characterization of synaptic properties near cell resting voltage. The voltage clamp measurements employed holding voltages of −70 mV and 0 mV to characterize excitatory and inhibitory synapses, respectively. For the time course characterization, we computed time to peak (time between the presynaptic spike and the peak), rise time (time between 20% and 80% of the peak at the rise stage), time of decay (the time constant from an exponential fit to the time course of decay, from 80% to 20% of the peak value), and the width (width at the half-peak level).

Results of this analysis indicate that the characteristics of all PSPs and PSCs are, on average, very similar between all the models (**Fig. S2**). The latter observation for the peak values is interesting, as the models were optimized separately based on overall characterization of network dynamics, and in principle one could expect bigger differences in synaptic weights (and, thus, PSP/PSC peaks) for the LL, LR, RL, and RR models. Furthermore, characteristics of PSPs and PSCs in the models are broadly consistent with the values reported in the literature, typically within a factor of 2 or less (e.g., [45, 47]; one should also note differences between the cortical regions and layers, different preparations and experimental conditions, as well as the naturally wide range of observed values). The trends observed in the literature were reproduced as well, such as faster dynamics of postsynaptic events in PV cells than in excitatory cells and a higher amplitude of inhibitory synaptic currents (e.g., [45]).

#### LGN filters

To enable simulations with arbitrary movies as visual stimuli, we developed a filter layer, the output of which was used to drive the L4 model, representing inputs from the lateral geniculate nucleus (LGN). The filters accepted movies (x,y,t-arrays) as inputs and produced time series of a firing rate as output. The output firing rates were taken to represent the rates of LGN neurons, which project to L4.

We used 3,000 filters each for three filter types – transient ON, OFF, and ON/OFF [6], i.e., representing 9,000 LGN cells. The transient types correspond to cells that produce an increase in firing rate to transitions from dark to bright (ON), bright to dark (OFF), or both (ON/OFF), such that the firing rate returns relatively quickly back to baseline levels after the change in scene. For simplicity, the sustained LGN cell types [6], for which the firing rate may differ substantially from the baseline after the initial transient (depending on the resulting brightness level), were neglected. The major reason for neglecting these types and overall representing the LGN inputs with rather simple filters was that little is known about how different LGN cell types combine their projections onto L4 neurons, and how properties of these projections are correlated with functional properties of the target cells.

Each LGN filter was represented by one (for ON and OFF filters) or two (for ON/OFF filters) receptive subfields, which were described by the following equations (see **Fig. 1**). The linear response, *L(t),* of a receptive subfield was given by

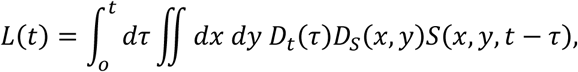

where *x* and *y* are the coordinates in the visual space (angles), *t* is the time, *S*(*x, y, t*) is the input signal (a movie), *D*_*t*_(*t*) is the temporal kernel, and *D*_*s*_(*x,y*) is the spatial kernel. For all operations with filters, we used a linear angle approximation for *x* and *y* (that is, replacing the tangent of an angle by the angle value in radians, which is approximately correct for small angles). The double-gaussian spatial kernel (center-surround) was used,

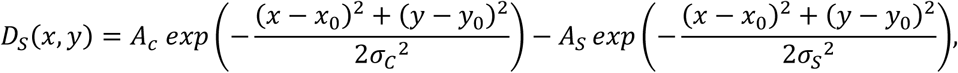

where we used *A*_*s*_ = *A*_*c*_/6, *σ*_*s*_ = 2*σ*_*c*_. The temporal kernel had the form

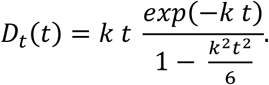

The input signal was grayscale, represented on a 0 to 255 scale (from black to white), with a time step of 1 ms and the pixel size of 1.25° in x and y (the frames were 192×96 pixels). It was fed directly into ON-type receptive subfields, or converted to 255 – *S*(*x,y, t*) and then fed into OFF-type receptive subfields. The linear response *L*(*t*) was then combined with the baseline rate *R*_0_ and passed through a rectifying nonlinearity, resulting in the firing rate:

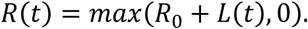

For the ON/OFF filter, *R*(*t*) from the ON and OFF receptive subfields were summed, and the mean of the corresponding *R*_0_ values was subtracted, to produce the final *R*(*t*) output.

The resulting firing rate *R*(*t*) was converted to spike trains using a Poisson random process, independently for each LGN filter (**Fig. 7a**). The firing rate exhibited a transient at the beginning of each visual stimulus, due to transition from no visual stimulation to a movie. To avoid this artefact, we always prepended 500 ms of full-field gray screen to a movie used as a visual stimulus, and replaced the resulting first 500 ms of *R*(*t*) simply by *R*_0_.

The centers of the individual filters, (*x*_0_, *y*_0_), were distributed randomly in the visual space, limited to approximately 130° x 90°. The two receptive subfields of each ON/OFF LGN cell were displaced with respect to each other (in random direction in x, y for each ON/OF cell), so as to enable orientation selectivity. Aside from that, the filters were axially symmetric, resulting in no orientation or direction selectivity. The temporal and spatial frequency (TF and SF) selectivity was thus primarily determined by the constants *σ*_*c*_, *σ*_*s*_, and *k* in the spatial and temporal kernels.

A new set of filters has been created for each of the three instantiations of the LL model type. These three instantiations of filter sets were then used for the three models within the LR, RL, and RR types.

The values of parameters for the filters were chosen randomly with a uniform probability within certain range (**Table S1**). The ranges were selected to produce approximately the same preferred SF and TF (0.05 cycles per degree, or cpd, and 4 Hz, respectively) across filters, which is approximately in the middle of the typical preferred SF and TF range in real LGN cells [6]. Although real LGN cells exhibit a wide range of preferred SF and TF values, we used the above simplification in order to avoid complexities of how different information processing channels with different SF and TF tuning converge on the L4 cells, as these aspects of feedforward connectivity from LGN to V1 are largely unknown. The specific choice of parameters also allowed for relatively wide tuning over SF and TF for our LGN filters, even though the peak was almost always at SF=0.05 cpd and TF=4 Hz. This is generally consistent with experimental observations, where most LGN cells exhibit smooth progression of diminishing responses away from preferred SF and TF, rather than very sharply tuned responses [6].

The above requirements were used to select values of *σ*_*c*_ and *k*. The baseline firing rate, *R*_0_, was selected to conform approximately to the levels of spontaneous activity exhibited by LGN cells [6] (**Table S1**). The remaining free parameter, *A*_*c*_, was selected so that the maximal observed F0 component (see below) of the responses to gratings was 11-12 Hz (**Table S1**), also approximating experimental findings [6]; +/−10% variation was allowed for this parameter.

It should be noted that for this parameterization and for analysis described below we used the definitions of the F0 and F1 components of responses to drifting gratings following Ref. [6]. Specifically, we used the cycle-averaged firing histograms for each cell (using the cycle period of the drifting grating stimulus); F0 was the mean rate computed from the histogram and F1 was the absolute value of the Fourier component at the frequency of the drifting grating stimulus (i.e., for a sine function, *a* + *b sin*(*ωt*), *F*_0_ = *a* and *F*_1_ = |*b*|).

Response properties of instantiated filters were analyzed and are presented in **Table S1**. Example responses of a transient ON and transient ON/OFF filters to gratings, in terms of tuning to orientation, SF, and TF, are also shown in **Fig. 7b** (responses of transient OFF cells are very similar to those of transient ON cells, except being inverted with respect to dark vs. bright). These analyses show that indeed the filters are not selective to the orientation and direction (except for F1 responses for transient ON/OFF cells), and are tuned almost exclusively to SF=0.05 cycles per degree (cpd) and TF=4 Hz. However, the SF and TF tuning of individual cells (**Fig. 7b**) is relatively broad, as intended.

#### Supplying inputs from LGN filters to L4 neurons

Feedforward connections from groups of LGN filters to the L4 cells were created based on the shared retinotopy. Whereas the LGN filters were defined in the visual space, the L4 cells were not, and for connecting them we introduced retinotopy in the L4 model by mapping the x, y coordinates (i.e., the plane of the layer) in the space of L4 cells to the x, y positions in the visual space. The center of the L4 model was assumed to correspond to the center of the visual space (i.e., center of the eye field). The positions were mapped using the conversion factor (from physical to visual space) of 120°/mm in x and 50°/mm in y, following the experimental observation that representation of azimuth and elevation on the surface of mouse V1 differs in length scale approximately by a factor of 2 [61]. With this mapping, each L4 cell was assigned an x,y position in the visual space.

To connect LGN cells to L4 cells, we created separate “lasso” subfields for each of three LGN types (transient ON, OFF, and ON/OFF), independently for each L4 cell (**Fig. 7c**; see also **Fig. 1** in Main Text). As above, a linear angle approximation was used here. The three subfields were positioned around the visual-space x, y location of the L4 cell. The lasso subfields for inhibitory L4 target cells were bigger, more symmetric, and more overlapping than those for excitatory target cells, since experimental studies show that receptive fields of fast-spiking interneurons tend to be larger than those of excitatory cells in L4 and L2/3, and orientation selectivity of fast-spiking interneurons is also significantly lower (see, e.g., [62]). Specifically, for target L4 cells of PV1, PV2, and inhibitory LIF types, all lasso subfields were centered at the L4 cell’s position and were circular, with the diameter randomly selected from the range [15°, 20°]. For the excitatory target cells, the ON/OFF lasso subfield was centered at the L4 cell’s position and was circular, whereas centers of the ON and OFF subfields were equidistant from the center, separated by an offset distance chosen randomly from the range of [10°, 11°] (thus enabling orientation selectivity and preference for a particular SF). These subfields were ellipsoidal (of equal size for a given target cell), with the ellipse aspect ratio being selected randomly from a range of [2.8, 3.0]. The ellipse minor radius was chosen randomly from the range [3°, 4°], and the major radius was computed by multiplying the minor radius by the aspect ratio. The minor axis was oriented along the line connecting the center of the two subfields. The ON/OFF subfield radius was equal to the minor radius of these ellipses.

For the excitatory L4 target cells, the choice of the direction in which the ON and OFF lasso subfields were positioned (**Fig. 7c**) was determined by the assigned preferred orientation angle that was set for each L4 cell at model construction. The angle between the vector connecting the centers of the ON and OFF subfields and the (1, 0) vector in the visual space was chosen to coincide with this assigned preferred orientation angle. Note that the actual preferred orientation angle was determined by the network activity in the simulations, and generally speaking did differ (more or less) from the assigned one.

Once all the lasso subfields were established for an L4 cell, the corresponding LGN cells were selected based on whether centers of their spatial kernels happened to be inside the subfields or not. Among those inside, a random set of LGN cells was selected, with the total number of each LGN type selected for one L4 target cell being capped at 15 for the inhibitory target cells and at 8 for the excitatory target cells (i.e., because we used 3 types of LGN cells, the maximum number of source LGN cells was 45 for inhibitory targets and 24 for excitatory targets). For the ON/OFF filters, the additional condition applied during the selection of sources, which required that selected filters were oriented (i.e., the axis connecting the centers of their ON and OFF subfield was oriented) within +/−15° from the assigned preferred orientation angle of the L4 cell. In terms of connections from individual LGN filters to L4 cells, after instantiating models we found broad diversity in the number of target L4 cells per LGN filter, from 0 to several hundred; on average, one LGN filter connected to 100 +/−170 L4 cells.

Since the LGN filters used did not represent all the LGN cell types (specifically, no sustained types [6]), we assumed that the number of synapses they provided to the L4 cells was smaller than the total expected number of synapses from the LGN (∼1,000, see Main Text). We thus set the number of synapses per each LGN filter connected to an L4 cell to be 30, resulting in approximately 600 synapses from the LGN for each L4 excitatory cell.

The geometric constraints above (size and separation of the lasso fields, etc.) were chosen to reflect approximately the characteristics of the receptive fields of the LGN inputs to L4 excitatory cells, based on *in vivo* measurements of LGN-only currents into L4 cells, resulting from visual stimulation [7], and also taking into account typical sizes of the receptive fields of LGN cells themselves, based on extracellular electrophysiological measurements [6]. Arguably, the constraints we applied were representative of a typical case, but did not reflect the considerable diversity of the receptive field sizes and shapes [6, 7]. This was done for simplicity, as we aimed to represent well the responsiveness of the LGN and L4 cells to SFs and TFs that are in the middle of typical values that evoke robust responses *in vivo* (SF∼0.05 cpd and TF∼4Hz). Introducing more diversity would require further assumptions about connectivity from LGN to L4 and within L4, in absence of data. Another note is that the choice of the geometry of “lasso” subfields (high aspect ratios) may appear to contradict observations of the rather symmetrical and highly overlapping subfields for LGN inputs to L4 [7]. However, due to the large size of individual LGN filter receptive fields, the combined receptive subfields for the LGN inputs to L4 cells were not extremely elongated and exhibited certain degree of overlap (see example on the right in **Fig. 7c**), similar to the experimental observations.

#### Stimulation protocol

All simulations were performed with visual stimuli chosen among drifting gratings, natural movies, natural images, moving bars, and full-field flashes (**Fig. S3**, the first three are part of the stimulus set in the Allen Brain Observatory [53]). Separate simulations with uniformly gray screen were also carried out to characterize spontaneous activity. The list of all visual stimuli used for all model systems is presented in **Table S2**. All stimuli were mapped (linearly subsampled or interpolated) to 192×96 pixels and time step of 1 ms. Each trial was a separate simulation. Typically, 10 trials were simulated for each stimulus condition, with some exceptions specified below.

Drifting gratings were square-wave alternating white and black stripes (typically at contrast 80%, although simulations with other contrasts were performed as well), drifting in the direction perpendicular to the stripes, at 0, 45, 90, 135, 180, 225, 270, or 315 degrees. For LGN filter characterization, 240 different gratings were used, in combinations of the 8 directions listed above, 6 SF (0.025, 0.05, 0.1, 0.2, 0.4, and 0.8 cpd), and 5 TF (1, 2, 4, 8, and 15 Hz). Due to the high computational expense associated with running simulations of the L4 model, and because the LGN filters were tuned to respond preferentially to SF ∼ 0.05 cpd, for the full model simulations we used a subset of these 240 conditions. A single SF=0.05 cpd and a few TF (typically, 2, 4, and 8 Hz) were used, with all 8 directions (**Table S2**). After the initial 500 ms of the gray screen, the gratings were presented for 2500 ms.

The gratings were named from g1 to g240 in the following manner: starting from the minimal TF, SF, and direction angle for g1, the numbering increased first along the TF dimension, then along the SF dimension, and finally along the direction dimension. Thus, for example, g1, g2, g3, g4, and g5 were all at direction 0 degrees, SF=0.025 cpd, and TF varying from 1 to 15 Hz; or, gratings with a given SF and TF, such as SF = 0.05 cpd and TF = 4 Hz, had numbers that were always separated by 30 – g8, g38, g68, g98, g128, g158, g188, g218, for directions 0, 45, 90, 135, 180, 225, 270, and 315 degrees, respectively. (See *Supplementary Files* and **Table S2** for more details.)

Natural movies were 3 clips from the opening scene of *Touch of Evil* [63, 53]. After the initial 500 ms of the gray screen, the movies were presented for 4500 ms. The three clips were named “TouchOfEvil_frames_N1_to_N2”, where N1 and N2 were the first and last frame based on the sequence from the original movie clip (these frames were 33 ms each, as the movie was encoded at 30 Hz rate; the clips used for simulations were upsampled to 1 kHz rate). The N1 and N2 were 1530 and 1680 for the first clip, 3600 and 3750 for the second, and 5550 and 5700 for the third.

For natural images, we used 10 examples taken from the Berkeley Segmentation Dataset [64] and the van Hateren Natural Image Dataset [65] (see also [53]). Each trial consisted of a 3000 ms long simulation, with the first 500 ms being the gray screen, and the rest covered by presentation of the 10 images in a random sequence, for 250 ms each, without interruption. We carried out simulations of 100 such trials, each named “imseq_i”, where i was 0 to 99 (“imseq” standing for “image sequence”).

Moving bars were single white or black stripes on gray background, moving with constant velocity in a direction perpendicular to the stripe. Four conditions were assessed – a black or white vertical bar moving left to right, and a black or white horizontal bar moving bottom to top. The bars were approximately 4 degrees wide and moved with the velocity of 33.8 degrees per second. These simulations were named “Bbar_v50pixps_hor”, “Bbar_v50pixps_vert”, “Wbar_v50pixps_hor”, and “Wbar_v50pixps_vert”.

Two types of full-field flashes were used (named “flash_1” and “flash_2”). For type 1, the screen was gray from 0 to 1000 ms, white from 1000 to 2000 ms, and gray again from 2000 to 3000 ms. For type 2, the screen was gray from 0 to 600 ms, white from 600 to 650 ms, and gray again from 650 to 1500 ms.

Spontaneous activity was measured during 1000 ms long presentation of gray screen. Typically, 20 trials were used. These simulations were named “spont”.

All analysis generally disregarded the first 500 ms of each simulation, as an equilibration period (with gray screen presented as a visual stimulus during this time in all cases).

For computational efficiency, a core set of stimuli from those described above was used for all systems (three cases each of LL, LR, RL, and RR), whereas the rest were applied in a targeted manner, primarily for the LL system (**Table S2**). Out of those, the richest set of stimuli was applied to model 2 of the LL type (“LL2”).

The LGN filter and background source projections to the L4 cells were generated independently for the 3 models of the LL type and then reused for the 3 models each of the LR, RL, and RR types. The same procedure was applied to recurrent connections for LL and LR cases, as well as for RL and RR. Furthermore, each trial of visual stimulation was accompanied by an independently generated trial of the background activity. These were generated independently for the 3 models of the LL type, and then applied consistently to the 3 models of the LR, RL, and RR types. Thus, for example, the LGN and background feedforward connections, as well as LGN and background spike trains incoming to the L4 cells on a particular trial of a given visual stimulus, were exactly the same between the models LL2, LR2, RL2, and RR2, but different from, e.g., LL1 or RL3.

#### Modeling optogenetic perturbations

A number of simulations was performed to mimic the effects of optogenetic manipulations (see below for experimental details). The experimental optogenetic perturbation involved silencing of Scnn1a cells expressing ArchR through application of yellow light. To model such a perturbation, we injected hyperpolarizing current into somata of cells in our simulations (**Fig. S7**). We selected the cell type to be directly perturbed (typically Scnn1a), and a fraction of the cells in this cell type (all the cells in the cell type that were outside of this fraction did not receive current injections), and applied the current injections to those cells.

Since the light spot used in the experiments was ∼2 mm in diameter, in most simulations we applied perturbation over the whole area of our model (which was ∼1.7 mm in diameter), including the LIF neurons. In a subset of simulations LIF neurons were not affected (this difference did not qualitatively affect the observed trends). Delivering direct current injections to LIF neurons was not supported in the NEURON implementation we used, and thus we opted for adding inhibitory synapses to LIF neurons that were selected for optogentic perturbations. A single such synapse was added to each silenced LIF neuron, and received a Poisson spike train at 100 Hz frequency. The synaptic weight was calibrated to result in the reduction of the LIF neuron’s firing rate similar to that effected by a −100 pA current injection to the soma of biophysically detailed cells in a benchmark simulation. To represent different current injections, we linearly scaled the synaptic weights for these synapses on the LIF neurons. The LIF neurons were selected for receiving such perturbation based on the fraction of cells represented by the perturbed cell type and the fraction of cells selected for perturbation within the type. For example, Scnn1a constituted 43.5 % of all excitatory biophysical neurons; if we were applying optogenetic perturbation to 20% of Scnn1a biophysical cells, then we correspondingly applied perturbation via inhibitory synapses to 8.7 % of all LIF excitatory cells.

The modeled optogenetic manipulation was applied at the beginning of the simulation and was maintained throughout the simulated time. For majority of these simulations, the injection current levels were −30, −50, −100, and −150 pA, whereas the fractions of Scnn1a population receiving such injections were 20, 50, and 100% (see **Table S2**).

Optogenetic silencing of the LGN [8] (**Fig. 5d**) was modeled by eliminating all spikes coming from LGN filters starting at a desired time. For this study, such a time was always set to 1000 ms from the beginning of a simulation.

### Experimental Methods

All experiments, animal treatment and surgical protocols were carried out with authorization from the Institutional Animal care and Use Committee of the Allen Institute in accordance with the Public Health Service (PHS) Policy on Humane Care and Use of Laboratory Animals.

#### Experimental characterization of LGN-to-L4 synapses

The synapse counts were performed using disectors on a systematic random sampling scheme. This allows for unbiased sampling and it has been used in other cortical areas and animals models (e.g. [66, 39]). We cannot however identify the cell type of the post synaptic target and therefore we assume that thalamic neurons target cell types according to the available dendrite of each cell. Though exceptions to this rule have been found in other areas or animal models, the contribution of L5 and L6 apical dendrites to the overall dendritic length in L4 is expected to be smaller than the dendrite from L4 neurons and one expects that it will not change substantially the proportion of thalamic synapses onto L4 in ways that would affect the predictions of the model.

We labeled thalamic boutons with immunohistochemistry for VGLUT2, selected sampling sites using rare systematic random sampling and then used the physical disector method introduced by [67] to perform the synaptic counts.

##### Histology

Wild type mice were perfused with fixative (2% paraformaldehyde and 0.5% glutaraldehyde) and a series of 60 um coronal slices through primary visual cortex (V1) was taken, and alternating slices were processed for electron microscopy (EM) with immunohistochemistry, or processed with Nissl stain for light microscopy (LM).

Slices selected for immunohistochemistry were incubated in anti vesicular glutamate transporter 2 (VGLUT2) antibody. Nickel intensified diamino benzidin or silver intensified gold particles were used to visualize the labeled elements in the electron microscope (EM). Slices were then osmicated, dehydrated and embedded in Durcupan resin.

We used as a primary antibody anti-VGLUT2 from Millipore, catalog number MAB5504 (lot 2322521). The secondary antibody was biotinylated-goat-anti-mouse IgG, from vector BA-9200 (lot X0623). Anti-VGLUT2 primary antibodies label thalamic terminals in several species including the mouse visual, somatosensory and motor cortices [68, 39] and it had been validated in layer 4 of mouse visual cortex by co-labeling of axons that have been injected in the thalamus [68]. The distribution of labeled terminals in this study matches the one previously observed in the cited works.

##### Sampling

Slices, grids, sections and sampling sites were chosen using a systematic random sampling (SRS). A random number in the range of the number of slices was generated via computer, and used to select target coronal slices for the EM series. A general region of interest in V1 was then selected based on comparison to the cytoarchitecture of the immediately adjacent Nissl-stained sections. This region of interest, which contained layer 4, was excised from the EM slice, trimmed, and sectioned with an ultramicrotome at 40 nm thickness and serial section ribbons collected onto copper grids. Labeled terminals were then examined and photographed in a Jeol 1200 EX II electron microscope using a digital camera (Gatan).

Low magnification EM images were overlaid and used to co-register the EM sections with the LM (Nissl) images, using vasculature and cell body (soma) landmarks. The Layer 4 region was determined by cyto-architecture, then drawn on the EM image.

##### Physical disector

A regular sampling grid of 45 × 45 μm was generated within this layer 4 region, and high magnification images were taken at each of these sampling points for disector analysis. Reference and lookup section were separated by one intervening ultrathin section so that the disector ‘*z*’ dimension was 0.08 μm. Both sections were used as reference and as lookup in order to double the sample [66, 69]. Three hundred and forty one disectors were used from 3 mice. The disectors had a size of 5 × 5 μm and at 3.6 nm per pixel. Synapses and associated structures were classified using conventional criteria [70, 71]. The percentage of thalamic synapses was calculated by dividing the number of synapses formed by labeled boutons by the total number of asymmetric synapses (putative excitatory).

#### Extracellular electrophysiological recordings in vivo

All electrophysiological recordings were performed in the left hemisphere of awake adult C57Bl/6 mice (2 to 6 months, males). A few weeks before the recordings, mice were first implanted with a metallic headplate using aseptic conditions and under anesthesia (see details in [6]). The day before or of recording, we performed a craniotomy over V1 (∼0.5 × 0.5 mm above the monocular portion of V1, 2.5 mm lateral to lambda) and implanted a reference skull screw; if the day before, then the exposed skull was sealed with Kwik Cast. On the day of recording the exposed cortex and skull were covered with 1% agarose in saline in order to prevent drying and to help maintaining mechanical stability. Dexamethasone was given to the animals to avoid brain inflammation (2 mg/kg, sc) and atropine to keep the respiratory tract clear (0.3 mg/kg, ip). Then, awake mice went into the recording setup. The headplate was clamped for stability, while the animal was free to run or remain still on a freely rotating disk. Multi-site silicon electrode arrays (imec Neuropixels) were dipped in Dil allowing post hoc visualization of the electrode path and lowered with a microdrive to a depth of 800-1000 μm. Reference and ground of the electrode were made through screws implanted in the skull. To improve the stability of recorded units, we allowed 20 minutes for the electrodes to settle.

Neurophysiological signals were amplified, band-pass filtered, and acquired continuously at 20 or 30 kHz at 16-bit resolution using an Amplipex system (Amplipex Ltd) or Open Ephys system (open-ephys.org). The spike sorting procedure was described in detail previously [72]. In brief, algorithmic identification and unit assignment of spikes [73, 74] was followed by manual adjustment of the clusters [75, 76]. Only units with clear refractory periods and well-defined cluster boundaries were included in the analyses [73].

Visual stimuli were largely the same as described above for simulations. The number of trials in the experiments was 50 for flashes, 10 for natural movies, and 100 for natural images. Rates of spontaneous activity were calculated by averaging the rates of each cell over 20 non-overlapping intervals, 500 ms long each, that were evenly distributed over 1 minute of continuous gray screen presentation (20 intervals of 500 ms were used for consistency with the 20 trials of gray screen presentation in simulations; see above). Moving bars were not used. Data for drifting gratings was taken from a previously published dataset [6].

After the recording, mice were perfused and brains fixed, sectioned, and stained with DAPI (see details of histology and imaging in [6]) for reconstruction of the electrode path to assign a layer to each recording site.

The laminar location of neurons was identified using histology and confirmed, when possible, by investigating the CSD depth profile of responses to full-field flashes. Putative excitatory neurons belonging to L4 according to this annotation were used for analysis. Since L4 is thin, and the fraction of inhibitory neurons is only ∼15%, we could not obtain a sizeable number of such neurons for L4 in our recordings. Because of that, putative fast-spiking inhibitory neurons from all layers were combined for analysis.

#### Whole-cell patch-clamp in vivo recordings

Data of *in vivo* intracellular membrane potential (*V*_*m*_) was obtained from genetically defined neuron populations in cortical layer 2/3 of mouse V1 via two-photon targeted whole-cell recordings. Adult (2 – 4 months old, n = 2) transgenic mice expressing a red fluorescent cytosol marker tdTomato (tdT) under the control of the Cux2-CreERT2 promoter were used. Recordings were performed under Isoflurane (1-1.5% in O_2_) anesthesia with ∼7 MΩ glass micropipettes filled with K-Gluconate based internal solution (containing, in mM: potassium gluconate 125, NaCl 10, HEPES 20, Mg-ATP 3, Na-GTP 0.4, in ddH_2_O; 290 mOsm; pH 7.3; Alexa 488,50 μg/ml). Exposed V1 region was covered with Artificial Cerebrospinal Fluid (ACSF) containing (in mM: NaCl 126, KCl 2.5, NaH_2_PO_4_ 1.25, MgCl_2_ 1, NaHCO_3_ 26, glucose 10, CaCl_2_ 2, in ddH_2_O; 290 mOsm; pH 7.3). tdT expressing neurons within 100 μm – 300 μm underneath the pia surface were targeted and patched, as described previously [77, 78, 79].

Two-photon imaging was performed using a Sutter Moveable Objective Microscope (Sutter, CA USA) coupled with a tunable Ti:Sapphire femtosecond laser (Chameleon Ultra II, Coherent USA), controlled by the open-source software ScanImage 3.8 (Janelia Research Campus/ Vidrio Technologies), to visualize L2/3 neurons in V1. Neurons were imaged with a 40x water-immersion objective (LUMPLFLN 40XW, Olympus USA) at the excitation wavelength of 920 nm. Electrophysiology Signals were amplified with a Multiclamp 700B, digitized with a Digidata 1440B at 20kHz, using pClamp software (Molecular Devices) and stored on a PC (Dell, USA). Quality of the pipette was checked both electrically and optically. Pipette would be discarded if disproportional increase of pipette resistance (Rp) occurred, which indicates occlusion of pipette tip. Optically checked ejection of dye under two-photon imaging was also used to indicate whether the pipette was free of occlusion. If both the electrical and optical measures agreed the tip is clogged, the penetration would be terminated and the pipette was retracted and discarded.

To conduct *in vivo* whole-cell recordings, clean pipette was manually advanced towards the target neuron with a low positive pressure (∼20 mbar above ambient) under two-photon imaging with Rp constantly monitored. Pipette capacitance was compensated before patching the cell. Standard procedures were used to rapidly form a gigaseal [77, 78, 79]. After gigaseal formation, a brief negative pressure was used to rupture the membrane inside the tip to achieve the whole-cell configuration. *V*_*m*_ was recorded under current clamp mode with series resistance (Rseries) appropriately compensated. Spontaneous *V*_*m*_ activity was recorded for at least 1 – 2 mins. The recordings were not corrected for liquid junction potential.

#### Optogenetic manipulations and extracellular electrophysiological recordings in vivo

We performed in vivo multichannel silicon probe recordings from the primary visual cortex of mice as previously described [52]. Briefly, mice were anesthetized with a combination of 5 mg/kg chlorprothixene (intraperitoneal) and 1.2 g/kg urethane (intraperitoneal); during surgery mice were supplemented with 0.5-1.0% isoflurane. A head frame with an ∼2 mm central opening was mounted over V1 and the skull was thinned using a dental drill. PBS was then applied to the thinned skull and sharpened forceps were then used to remove a small portion of skull wide enough to permit insertion of a NeuroNexus 32-channel linear probe (A1×32-Edge-5mm-20-177). The probe was inserted at a depth of 800-1,000 μm. PBS was then applied to keep the craniotomy moist. Following a recovery period following probe insertion of at least 20 minutes, visual stimuli were presented using a gamma-corrected, Dell 52 3 32.5 cm LCD monitor (60 Hz refresh rate, mean luminance 50 cd/m2, 25 cm from contralateral eye). We used the Psychophysics Toolbox [80] to generate and present full-field sinusoidal drifting gratings (temporal frequency, 2 Hz; Spatial frequency, 0.04 cycles per degree; 2 orientations, 0 and 90 degrees; contrasts 0, 10, 18, 32, 64, and 100% contrast). Drifting gratings were presented for 1500 ms and optogenetic stimulation trials were interleaved with non-optogenetic trials. Each trial was separated with 3-6 s inter-trial interval with a gray screen. To photostimulate archaerhodopsin (optogenetic trials) we used a 1 mm LED fiber with ∼20 mW power output at the fiber tip, which was placed 0.5 mm from skull above recording location (590 nm, 1 mm diameter, Doric Lenses). The optogenetic light delivery lasted 1.9 s and started 200 ms prior to the visual stimulus and ended 200 ms following the stimulus. Black foil (Thor Labs) was shaped into a shield around LED and headframe in order to prevent the LED light from reaching the eyes. Recordings were amplified 1,000X and band-pass filtered between 0.3 Hz and 5 kHz using an AM systems 3500. Acquisition was done at 20 kHz with a NIDAQ PCIe-6239 board using custom MATLAB software (MathWorks).

Data were analyzed with custom-written software using MATLAB. Single units were isolated using software provided by D.N. Hill, S.B. Mehta, and D. Kleinfeld [81]. Signals were high-pass filtered at 500 Hz and waveforms were extracted from four adjacent electrode sites. Spikes were defined as events exceeding 4-5 SD of the noise. Waveforms were clustered using a k-means algorithm and further aligned using a graphical user interface. Fisher linear discriminant analysis and refractory period violations were used to assess unit isolation quality. Units were assigned a depth based on the channel in which they showed the strongest signal. In this work, only the data corresponding to drifting gratings with 100% contrast was used.

The following mouse lines were used for in vivo extracellular recordings: Scnn1a-Tg3-Cre (stock number: 009613; B6;C3-Tg(Scnn1a-cre)3Aibs/J) and Ai35(RCL-Arch/GFP) (stock number: 012735; B6;129S-Gt(ROSA)26Sortm35.1(CAG-aop3/GFP)Hze/J).

### Data Analysis

Data analysis was performed on all simulations carried out for a given stimulus, as described in **Table S2**. That typically involved at least two independent models (usually three models) and multiple trials. Data from these multiple models and trials were typically combined together to produce summary plots.

Summary figures that present box plots of various analyzed quantities adhere to the following standard. The box bottom and top represent the lower boundary of the second quartile of the data and the top boundary of the third quartile, respectively, thus marking the inter-quartile range (IQR); the median is shown in red. Whiskers mark +/−1.5 IQR from the bottom and top of the plot, as long as there are data points in that range. The rest of the data (outliers) are shown as separate dots or other symbols. In cases when the plot range does not include the whole span of the data (which was sometimes necessary due to the log-normal-like distribution of the data points over several orders of magnitude), the outliers outside of the plot range are not shown explicitly but are instead indicated by an arrow. The numbers next to such arrows (“N1/N2”) indicate the number of such outliers that are not shown (N1) and the total number of data points (N2). The mean and s.e.m. of the data are shown on these plots as circles with error bars (thick black lines).

#### Computation of the features of firing rate responses

Spontaneous activity was characterized by analyzing the responses of cells from 20 gray screen trials, each 1,000 ms long. For each cell, the spikes from the second 500 ms of every trial were used to compute the firing rate, which was then averaged over trials.

The maximal response to gratings (*R*_*max*_), the orientation selectivity index (OSI) and the direction selectivity index (DSI) were computed using responses to drifting gratings (disregarding the first 500 ms of the gray screen). First, the firing rate for each cell was obtained, using spikes from the 2,500 ms of grating presentation, and averaging over all trials. This rate was computed separately for every grating condition. The *R*_*max*_ metric for each cell was the maximal firing rate among all conditions.

Following [6], we used the firing rate directly to compute the OSI and DSI (i.e., without subtracting the spontaneous rate). The analysis was performed for each cell independently. For each orientation, an average response across all SF and TF was computed, and the orientation that corresponded to the greatest averaged response was assigned as the preferred orientation of the cell. The SF and TF that evoked the strongest response at the preferred orientation were then considered as preferred frequencies. OSI and DSI were computed using the responses at all orientations for these preferred SF and TF. The OSI was computed as one minus the circular variance,

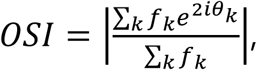

where *k* is the index for summation over all directions *θ*_*k*_, and *f*_*k*_ is the firing rate response for each direction. The DSI was computed as

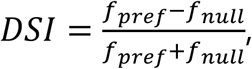

where *f*_*pref*_ is the firing rate response at the preferred direction (*θ*_*pref*_) and *f*_*null*_ is the response at the opposite direction (*θ*_*pref*_ + *π*).

Responses to flashes were characterized by averaging firing rates of multiple cells in multiple trials, in 2 ms bins. The number of trials was 10 for simulations and 50 for experiments. For simulations, the averaging was done separately for each of the two models for which 50 ms flash stimuli were simulated, and for each of the three biophysical excitatory cell populations, resulting in 6 firing rate vs. time traces. For experiments, the averaging was done separately for each mouse over all L4 excitatory cells, resulting in 8 traces. The peak magnitude and time to peak from the flash onset were then computed on each of those traces for the 1st and 2nd peaks.

#### Spectra of the local field potential (LFP) and of spiking activity

The extracellular potential Φ was calculated along the central axis of the model (perpendicular to the L4 plane), at locations distributed with the increments of 10 μm along the axis. The extracellular potential at the *j*-th location (i.e., electrode site) for a particular cell was computed as Φ_*j*_ = ∑_*k*_ *R*_*jk*_*I*_*k*_, where *I*_*k*_ is the membrane current through the *k-*th neuronal compartment and *R*_*jk*_ is the transfer resistance between *k-*th neuronal compartment and the *j*-th electrode site. The transfer resistances were computed using the line-source approximation [82] assuming homogeneous and isotropic extracellular conductivity of 0.3 S/m. The contributions of the individual cells to the recordings are then summed up to find the total recorded potential at each recording electrode site. The membrane currents were obtained with NEURON’s cvode.use_fast_imem() function which returns the transmembrane currents without needing to utilize the computationally expensive extracellular mechanism. The computation of the extracellular potential was implemented by modifying the NEURON’s advance() function—called on each time step—to include the function call to our Python module computing the extracellular potential.

The local field potential (LFP) was obtained by low-pass filtering the simulated extracellular potential with the 6th-order Butterworth filter, with the cutoff frequency of 200 Hz. The power spectra of the LFP (computed employing the fast Fourier transform) were then used for analysis of oscillations in population activity. The LFP recorded at the site closest to the center of the model was utilized for this analysis.

Power spectra of spiking activity (**Fig. S6**) were computed as follows. A position in space was selected for recording; in all cases in this work we chose the center of the model for this purpose. The signal was then computed as the inverse-distance weighted multiunit activity, i.e., the time-binned firing rate accumulated from all neurons with the weight 1/*r*_*i*_, where *r*_*i*_ is the distance from the neuron’s soma to the recording position. The time bin of 1 ms was used. The power spectrum was then computed using fast Fourier transform.

Both LFP and spiking activity spectra were averaged over all trials of a given stimulus.

#### Computation of sparsity

Sparsity of responses was computed following the definition in [50]. The formula used was

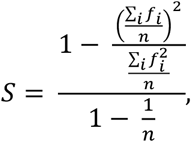

where *S* is the sparsity, *f* is the response (we used the firing rates averaged for each cell within each time bin over all trials for a given visual stimulus), and *n* is the number of samples *i*. The lifetime sparsity can be defined for each cell, and then samples *i* correspond to time bins distributed over the time axis, for a given visual stimulus.

For purposes of illustrating sparsity in our simulations, we computed lifetime sparsity for the three natural movies and three drifting grating stimuli (SF=0.05 cpd, TF=4 Hz, orientation of 0, 45, and 90 degrees). Since gratings were presented for 2.5 seconds vs. 4.5 seconds for the movies, only the first 2.5 seconds of movie responses were used for this analysis (to enable equal comparison).

#### Measurement of the excitatory currents in the model

To measure the currents flowing through the somata of biophysical cells, we used NEURON’s function SEClamp(), which provides a simple voltage clamp. Out of all 10,000 biophysical cells in a model, every 200th cell was voltage-clamped at the soma at −70 mV, to measure the excitatory current [7]. Thus, the currents were recorded for 50 cells in such a simulation (e.g., cell ID 2, 202, 402, 602, 802, …, 9802). The currents were saved from the SEClamp() objects, at each simulation time step.

For the measured current, the F0 (mean) and F1 (amplitude at the frequency of the stimulus) components were obtained according to Ref. [7]. The current was averaged over trials and then cycle-averaged. F0 was computed as the mean (over the cycle time) of the cycle-averaged current. F0 was then subtracted from the cycle-averaged current, and the result was fit with a sinusoid function at the stimulus frequency, *b* sin(*ωt* + *φ*), where *ω* = 2*πv* (*v* being the stimulus frequency). The F1 component value was taken as *F*_1_ =2|*b*|.

For all-LIF models, we measured contribution of LGN synaptic inputs to the total excitatory synaptic inputs as a proxy to the excitatory synaptic currents measured for the biophysical model. Specifically, for each spike arriving to a synapse, that synapse contributed its weight to the accumulating sum for the target cell; the sum was computed for synapses from LGN only or for all synapses and was normalized by time and number of trials.

